# A high resolution atlas of gene expression in the domestic sheep (*Ovis aries*)

**DOI:** 10.1101/132696

**Authors:** EL Clark, SJ Bush, MEB McCulloch, IL Farquhar, R Young, L Lefevre, C Pridans, HG Tsang, C Wu, C Afrasiabi, M Watson, CB Whitelaw, TC Freeman, KM Summers, AL Archibald, DA Hume

## Abstract

Sheep are a key source of meat, milk and fibre for the global livestock sector, and an important biomedical model. Global analysis of gene expression across multiple tissues has aided genome annotation and supported functional annotation of mammalian genes. We present a large-scale RNA-Seq dataset representing all the major organ systems from adult sheep and from several juvenile, neonatal and prenatal developmental time points. The *Ovis aries* reference genome (Oar v3.1) includes 27,504 genes (20,921 protein coding), of which 25,350 (19,921 protein coding) had detectable expression in at least one tissue in the sheep gene expression atlas dataset. Network-based cluster analysis of this dataset grouped genes according to their expression pattern. The principle of ‘guilt by association’ was used to infer the function of uncharacterised genes from their co-expression with genes of known function. We describe the overall transcriptional signatures present in the sheep gene expression atlas and assign those signatures, where possible, to specific cell populations or pathways. The findings are related to innate immunity by focusing on clusters with an immune signature, and to the advantages of cross-breeding by examining the patterns of genes exhibiting the greatest expression differences between purebred and crossbred animals. This high-resolution gene expression atlas for sheep is, to our knowledge, the largest transcriptomic dataset from any livestock species to date. It provides a resource to improve the annotation of the current reference genome for sheep, presenting a model transcriptome for ruminants and insight into gene, cell and tissue function at multiple developmental stages.

**Author Summary:** Sheep are ruminant mammals kept as livestock for the production of meat, milk and wool in agricultural industries across the globe. Genetic and genomic information can be used to improve production traits such as disease resiliance. The sheep genome is however missing important information relating to gene function and many genes, which may be important for productivity, have no informative gene name. This can be remedied using RNA-Sequencing to generate a global expression profile of all protein-coding genes, across multiple organ systems and developmental stages. Clustering genes based on their expression profile across tissues and cells allows us to assign function to those genes. If for example a gene with no informative gene name is expressed in macrophages and is found within a cluster of known macrophage related genes it is likely to be involved in macrophage function and play a role in innate immunity. This information improves the quality of the reference genome and provides insight into biological processes underlying the complex traits that influence the productivity of sheep and other livestock species.

## Introduction

Sheep (*Ovis aries*) represent an important livestock species globally and are a key source of animal products including meat, milk and fibre. They remain an essential part of the rural economy in many developed countries and are central to sustainable agriculture in developing countries. They are also an important source of greenhouse gases [1]. Although genetic improvement is often considered to have been less effective in sheep than in the dairy cattle, pig and poultry sectors, advanced genomics-enabled breeding schemes are being implemented in New Zealand and elsewhere [2-4]. A better understanding of functional sequences, including transcribed sequences and the transcriptional control of complex traits such as disease resilience, reproductive capacity, feed conversion efficiency and welfare, will enable further improvements in productivity with concomitant reductions in environmental impact.

RNA-Sequencing (RNA-Seq) has transformed the analysis of gene expression from the single-gene to the genome-wide scale allowing visualisation of the transcriptome and redefining how we view the transcriptional control of complex traits (reviewed in [5]). Large-scale gene expression atlas projects have defined the mammalian transcriptome in multiple species, initially using microarrays [6-9] and more recently by the sequencing of full length transcripts or of 5’ ends, for example in the horse [10], and in human and mouse by the FANTOM 5 consortium [11-13], ENCODE project [14] and Genotype-Tissue Expression (GTEx) Consortium [15].

These efforts have focused mainly on mice and humans, for which there are high quality richly annotated reference genome sequences available as a frame of reference for the identification and analysis of transcribed sequences. Draft reference genome sequences have been established for the major livestock species (chicken, pig, sheep, goat and cattle) over the past decade, yet it is only with the recent deployment of long read sequencing technology that the contiguity of the reference genome sequences for these species has improved. This is exemplified by the recent goat genome assembly [16, 17]. In these species there are still many predicted protein-coding and non-coding genes for which the gene model is incorrect or incomplete, or where there is no informative functional annotation. For example, in the current sheep reference genome, Oar v3.1 (Ensembl release 87) (http://www.ensembl.org/Ovis_aries/Info/Index), 30% of protein-coding genes are identified with an Ensembl placeholder ID [18]. Given the high proportion of such unannotated genes many are likely to be involved in important functions. Large-scale RNA-Seq gene expression datasets can be utilised to understand the underlying biology and annotate and assign function to such unannotated genes [19]. With sufficiently large datasets, genes form co-expression clusters, which can either be generic, associated with a given pathway or be cell-/tissue-specific. This information can then be used to associate a function with genes co-expressed in the same cluster, a logic known as the ‘guilt by association principle’ [20]. Detailed knowledge of the expression pattern can provide a valuable window on likely gene function, as demonstrated in pig [6], sheep [18, 21], human and mouse [8, 9, 22, 23].

A high quality well-annotated reference genome is an exceptionally valuable resource for any livestock species, providing a comparative sequence dataset and a representative set of gene models. The International Sheep Genomics Consortium (ISGC) released a high quality draft sheep genome sequence (Oar v3.1) in 2014 [18]. Included in the sheep genome paper were 83 RNA-Seq libraries from a Texel gestating adult female, 16 day embryo, 8 month old lamb and an adult ram. This Texel RNA-Seq transcriptome significantly improved the annotation of Oar v3.1 and identified numerous genes exhibiting changes in copy number and tissue specific expression [18]. To build on this resource and further improve the functional annotation of Oar v3.1 we have generated a much larger high-resolution transcriptional atlas from a comprehensive set of tissues and cell types from multiple individuals of an outbred cross of two economically important sheep breeds. To maximize heterozygosity we deliberately chose a cross of disparate breeds, the Texel, which is used as a terminal sire as it is highly muscled in some cases because it has a myostatin variant linked to double-muscling and meat quality [24], and the Scottish Blackface, a breed selected for robustness on marginal upland grazing [25].

The sheep gene expression atlas dataset presented here is the largest of its kind from any livestock species to date and includes RNA-Seq libraries from tissues and cells representing all the major organ systems from adult sheep and from several juvenile, neonatal and prenatal developmental time points. Because the tissues were obtained from multiple healthy young adult animals, the atlas may also aid understanding of the function of orthologous human genes. Our aim was to provide a model transcriptome for ruminants and give insight into gene, cell and tissue function and the molecular basis of complex traits. To illustrate the value of the resource, we provide detailed examination of genes implicated in innate immunity and the advantages of cross breeding and provide putative gene names for hundreds of the unannotated genes in Oar v3.1. The entire data set is available in a number of formats to support the research community and will contribute to the Functional Annotation of Animal Genomes (FAANG) project [26, 27].

## Results and Discussion

### Scope of Sheep Gene Expression Atlas Dataset

This sheep gene expression atlas dataset expands on the RNA-Seq datasets already available for sheep, merging a new set of 429 RNA-Seq libraries from the Scottish Blackface x Texel cross (BFxT) with 83 existing libraries from Texel [18]. Details of the new BFxT libraries generated for the sheep gene expression atlas, including the developmental stages sampled, tissue/cell types and sex of the animals are summarised in Table 1. These samples can be grouped into 4 subsets (“Core Atlas”, “GI Tract Time Series”, “Early Development” and “Maternal Reproductive Time Series”). The animals used to generate the four subsets of samples are detailed in S1 Table.

**Table 1:** Details of the tissues and cell types sequenced to generate the BFxT RNA-Seq dataset for the sheep gene expression atlas. Tissues and cells were chosen to cover all major organ systems. All libraries were Illumina 125bp paired end stranded libraries. See S2 Table for a detailed list of the tissues and cell types sequenced.

| Subset | Tissue Type | Library Type | Sequencing Depth | Total Number of Libraries | Number of Individuals |
| --- | --- | --- | --- | --- | --- |
| Core Atlas | Liver, spleen, ovary, testes, | Total RNA-Seq | >100 million reads per | 60 (10 per individual) | 3 adult males and 3 adult |
|  | hippocampus,<br>kidney medulla,<br>bicep muscle,<br>reticulum,<br>ileum, thymus,<br>left ventricle |  | sample |  | females |
| Core Atlas | GI tract,<br>reproductive<br>tract, brain,<br>endocrine,<br>cardiovascular,<br>lymphatic,<br>musculo-<br>skeletal,<br>immune cells | mRNA-Seq | >25 million<br>reads per<br>sample | 248 (~45 per<br>individual) | 3 adult males<br>and 3 adult<br>females |
| LPS Time<br>Course<br>(Core Atlas) | BMDM 0h (-<br>LPS) and 7h<br>(+LPS) | Total RNA-Seq | >100 million<br>reads per<br>sample | 12 (2 time<br>points per<br>individual) | 3 adult males<br>and 3 adult<br>females |
| LPS Time<br>Course<br>(Core Atlas) | BMDM 2h, 4h,<br>24h post LPS<br>treatment | mRNA-Seq | >25 million<br>reads per<br>sample | 32 (3 time<br>points per<br>individual) | 3 adult males<br>and 3 adult<br>females |
| GI Tract Time<br>Series | Gastro-intestinal<br>tract | mRNA-Seq | >25 million<br>reads per<br>sample | 90 (10 tissues<br>per individual) | 9 lambs (3 at<br>birth, 3 at one<br>week and 3 at<br>8 weeks) |
| Early<br>Development<br>(blastocysts) | Day 7<br>blastocysts (3<br>pools of 8) | Nu Gen Single<br>Cell Ovation<br>Kit | >66 million<br>reads per<br>sample | 3 | 3 pools |
| Early<br>Development<br>(day 23) | Day 23 whole<br>embryos | Total RNA-Seq | >100 million<br>reads per<br>sample | 3 | 3 |
| Early<br>Development<br>(day 35) | Liver, brain,<br>embryonic<br>fibroblasts | mRNA-Seq | >25 million<br>reads per<br>sample | 7 | 3 embryos<br>(two female<br>and one male) |
| Early<br>Development<br>(day 100) | Liver, ovary | mRNA-Seq | >25 million<br>reads per<br>sample | 5 | 3 female<br>(liver)<br>2 female<br>(ovary) |
| Maternal<br>Reproductive<br>Time Series<br>(days, 23, 35<br>and 100) | Placenta and<br>ovary | mRNA-Seq | >25 million<br>reads per<br>sample | 12 | 2 females per<br>time point |

The “Core Atlas” subset was generated using six adult virgin sheep, approximately 2 years of age. Tissue samples were collected from all major organ systems from 3 males and 3 females to ensure, wherever possible, there were biological replicates from each sex to support an analysis of sex-specific gene expression. In addition, five cell types were sampled, including peripheral blood mononuclear cells (PBMCs) and blood leukocytes. Since macrophages are known to be a highly complex source of novel mRNAs [28], and were not sampled previously, three types of macrophage (+/- stimulation with lipopolysaccharide (LPS)) were included.

For the “GI Tract Time Series” subset of samples we focused on 10 regions of the gastro-intestinal (GI) tract, immediately at birth prior to first feed, at one week and at 8 weeks of age. These time points aimed to capture the transition from milk-feeding to rumination. Embryonic time points were chosen, at days 23, 35 and 100, to detect transcription in the liver, ovary and brain in “Early Development”. Parallel time points were included for placenta and ovary samples from gestating BFxT ewes, comprising the “Maternal Reproductive Time Series” subset. Finally, 3 pools of eight day 7 blastocysts were included to measure transcription pre-implantation and these were also included in the “Early Development” subset.

A detailed list of all tissues and cell types included in each subset of samples can be found in S2 Table. Tissues and cell types were chosen to give as comprehensive a set of organ systems as possible and include those tissues relevant for phenotypic traits such as muscle growth and innate immunity.

### Sequencing Depth and Coverage

Approximately 37x10^9^ sequenced reads were generated from the BFxT libraries, generating approximately 26x10^9^ alignments in total. The raw number of reads and percentage of alignable reads per sample are included in S3 Table. For each tissue a set of expression estimates, as transcripts per million (TPM), were obtained using the high speed transcript quantification tool Kallisto [29]. Kallisto is a new transcriptome-based quantification tool that avoids the considerable bias introduced by the genome alignment step [30]. Gene level expression atlases are available as S1 Dataset and, with expression estimates averaged per tissue per developmental stage, S2 Dataset. The data were corrected for library type (as we described in [31] and summarised in S1 Methods). We used Principal Component Analysis (PCA) pre- and post-correction (S1 Fig) for library type to ensure the correction was satisfactory. Hierarchical clustering of the samples is included in (Fig 1) and illustrates both the large diversity and logical clustering of samples of samples included in the dataset.

**Fig 1:**
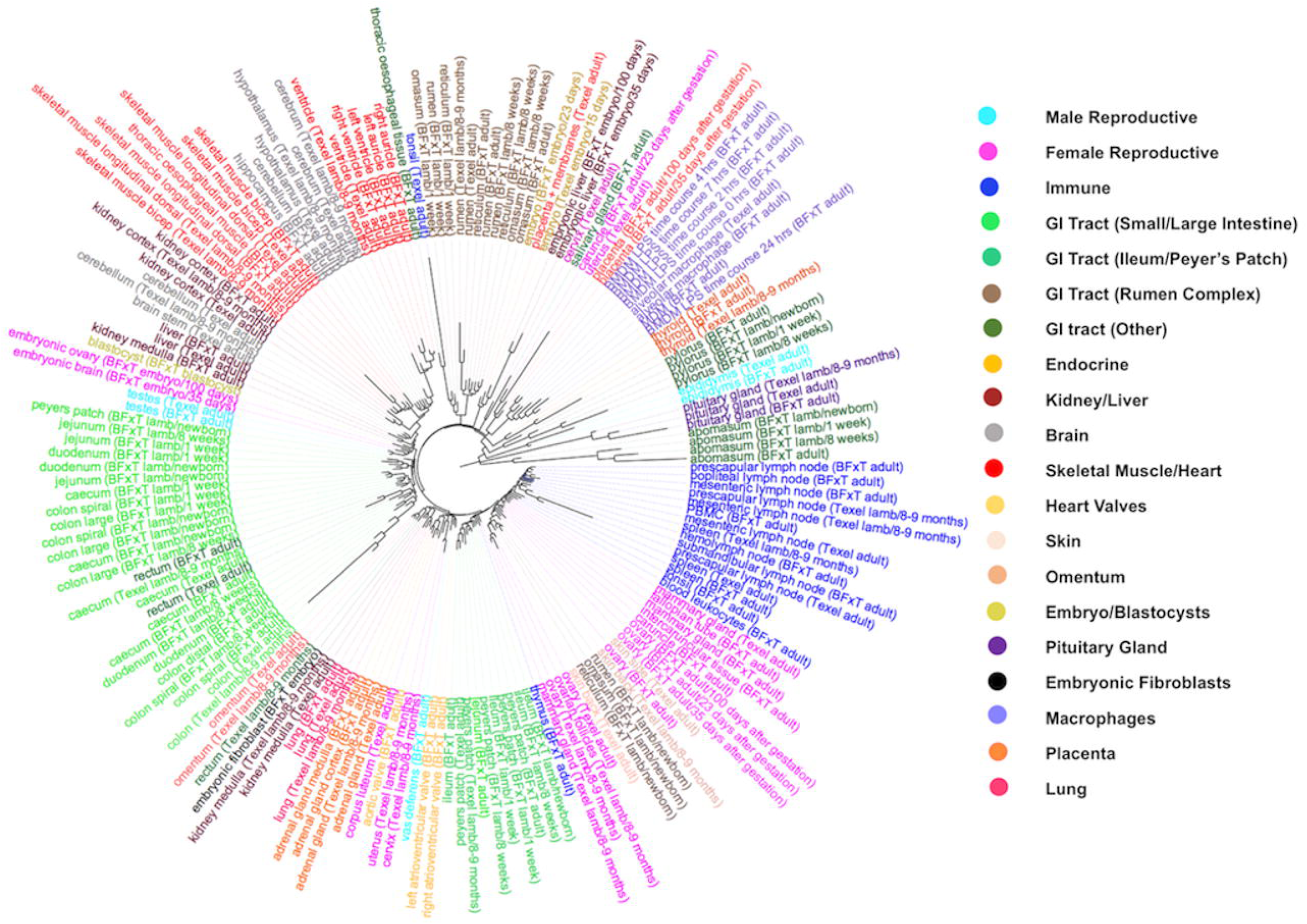
Hierarchical clustering of the samples included in the sheep gene expression atlas dataset. Samples of each tissue and cell type from each breed and developmental stage were averaged across individuals for ease of visualisation. The tree was constructed from the Euclidean distances between expression vectors using MEGA v7.0.14 [141] with the neigh-bourjoining method and edited in the graphical viewer FigTree v1.4.3 [142]. Clustering is biologically meaningful and highlights the lack of any significant effect of library type post-correction. Samples are coloured by organ system.

The *O. aries* reference genome (Oar v3.1) includes 27,504 loci that are transcribed (20,921 protein coding), of which 25,350 (19,921 protein coding) (97%) were detectable with expression of TPM >1, in at least one tissue from at least one individual, in the sheep gene expression atlas dataset, demonstrating the depth and scope of this dataset. The proportion of transcripts with detectable expression, after each ‘pass’ with Kallisto (See Materials and Methods), is presented in Table 2. Only 3% (561) of transcripts from Oar v3.1 (S4 Table) did not meet the minimum detection threshold of TPM > 1 in at least one tissue and therefore were not detected in the sheep atlas dataset. In a minority of cases, transcripts were missing because they were highly specific to a tissue or cell which was not sampled, such as odontoblasts (which uniquely produce tooth dentin, mediated by DSPP (dentin sialophosphoprotein) [32]). We also did not include any samples taken from the eye which expresses multiple unique proteins e.g., the lens-specific enzyme LGSN (lengsin) [33]. The majority (77%) of the genes not detected in the sheep atlas were unannotated, with no assigned gene name. A small number of these genes (36) lack sequence conservation and coding potential and so are potentially spurious models (S4 Table).

**Table 2:** The number and percentage of Oar v3.1 protein coding and non-coding genes, with average TPM across all animals > 1 in at least one tissue, in both the BFxT dataset after the Kallisto first and second pass, and after incorporating the existing Texel dataset. ‘BFxT data’ refers to the present study; ‘Texel data’ is obtained from [18]. The ‘first pass’ Kallisto index contains the known *Ovis aries* v3.1 cDNAs for both protein-coding and non-protein coding transcripts. The ‘second pass’ Kallisto index is a filtered version of the former, that (a) restricts the RNA space to protein-coding genes, pseudogenes, and processed pseudogenes (so that expression within an equivalent space will be quantified, irrespective of experimental protocol), (b) omits genes that had no detectable expression across all BFxT samples, and (c) includes novel transcript reconstructions further to the *de novo* assembly of unmapped reads.

| RNA-Seq data used: | Scottish Blackface x Texel Libraries Only |  |  |  |  | Including the existing Texel dataset |  |
| --- | --- | --- | --- | --- | --- | --- | --- |
| Kallisto index: | First-Pass |  |  | Second-Pass (restricted) |  | Second-Pass (restricted) |  |
| Gene type | No. in reference annotation (Oar v3.1) | No. of genes of this type expressed | % genes of this type expressed | No. of genes of this type expressed | % genes of this type expressed | No. of genes of this type expressed | % genes of this type expressed |
| lincRNA | 1858 | 1548 | 83.32 | 0 | 0 | 0 | 0 |
| miRNA | 1305 | 1242 | 95.17 | 0 | 0 | 0 | 0 |
| misc RNA | 361 | 310 | 85.87 | 0 | 0 | 0 | 0 |
| MT rRNA | 2 | 2 | 100 | 0 | 0 | 0 | 0 |
| MT tRNA | 22 | 19 | 86.36 | 0 | 0 | 0 | 0 |
| processed pseudogene | 43 | 31 | 72.09 | 35 | 81.40 | 38 | 88.37 |
| protein-coding | 20921 | 19921 | 95.22 | 20189 | 96.50 | 20359 | 97.31 |
| pseudogene | 247 | 172 | 69.64 | 189 | 76.52 | 201 | 81.38 |
| rRNA | 305 | 272 | 89.18 | 0 | 0 | 0 | 0 |
| snoRNA | 756 | 717 | 94.84 | 0 | 0 | 0 | 0 |
| snRNA | 1234 | 1116 | 90.44 | 0 | 0 | 0 | 0 |
| **Sum** | **27054** | **25350** |  | **20413** |  | **20598** |  |

### Gene Annotation

In the Oar v3.1 annotation, 6217 (~30%) of the protein coding genes lack an informative gene name. Whilst the Ensembl annotation will often identify homologues of a sheep gene model, the automated annotation pipeline used is conservative in its assignment of gene names and symbols. Using an annotation pipeline (described in S1 Methods and illustrated in S5 Table) we were able to utilise the sheep gene expression atlas dataset to annotate >1000 of the previously unannotated protein coding genes in Oar v3.1 (S6 Table). These genes were annotated by reference to the NCBI non-redundant (nr) peptide database v77 [34] and assigned a quality category based on reciprocal percentage identity, if any, to one of 9 known ruminant proteomes (S7 Table). A short-list containing a conservative set of gene annotations, to HGNC (HUGO Gene Nomenclature Committee) gene symbols, is included in S8 Table. Many of these genes are found in syntenic regions, and are also supported by the up- and downstream conservation of genes in a related genome, cattle (*Bos taurus* annotation UMD 3.1). S9 Table contains the full list of genes annotated using this pipeline. Many unannotated genes can be associated with a gene description, but not necessarily an HGNC symbol; these are also listed in S10 Table. We manually validated the assigned gene names on this longer list using network cluster analysis and the “guilt by association” principle.

### Network Cluster Analysis

Network cluster analysis of the sheep gene expression atlas was performed using Miru (Kajeka Ltd, Edinburgh UK), a tool for the visualisation and analysis of network graphs from big data [35-37]. The the atlas of unaveraged TPM estimates, available as S1 Dataset, were used for the network cluster analysis. The three blastocyst samples were removed from the network cluster analysis as they were generated using a library preparation method which was not corrected for and created a significant effect of library type. With a Pearson correlation co-efficient threshold of *r*=0.75 and MCL (Markov Cluster Algorithm [38]) inflation value of 2.2, the gene-to-gene network comprised 15,129 nodes (transcripts) and 811,213 edges (correlations above the threshold value). This clustering excludes >30% of detected transcripts, most of which had idiosyncratic expression profiles. One of the major sources of unique expression patterns is the use of distinct promoters in different cell types. The transcription factor *MITF* (Melanogenesis Associated Transcription Factor), for example, does not cluster with any other transcripts in sheep and is known in humans to have at least 7 distinct tissue-specific promoters in different cell types, including macrophages, melanocytes, kidney, heart and retinal pigment epithelium [13].

The resultant correlation network (Fig 2) was very large and highly structured comprising of 309 clusters ranging in size. Genes found in each cluster are listed in S11 Table and clusters 1 to 50 (numbered in order of size; cluster 1 being made up of 1199 genes) were annotated by hand and assigned a broad functional ‘class’ and ‘sub-class’ (Table 3). Functional classes were assigned based on GO term enrichment [39] for molecular function and biological process (S12 Table) and gene expression pattern, as well as comparison with functional groupings observed in the pig expression atlas [6]. Fig 3 shows a network graph with the nodes collapsed, and the largest clusters numbered 1 to 30, to illustrate the relative number of genes in each cluster and their functional class.

**Fig 2:**
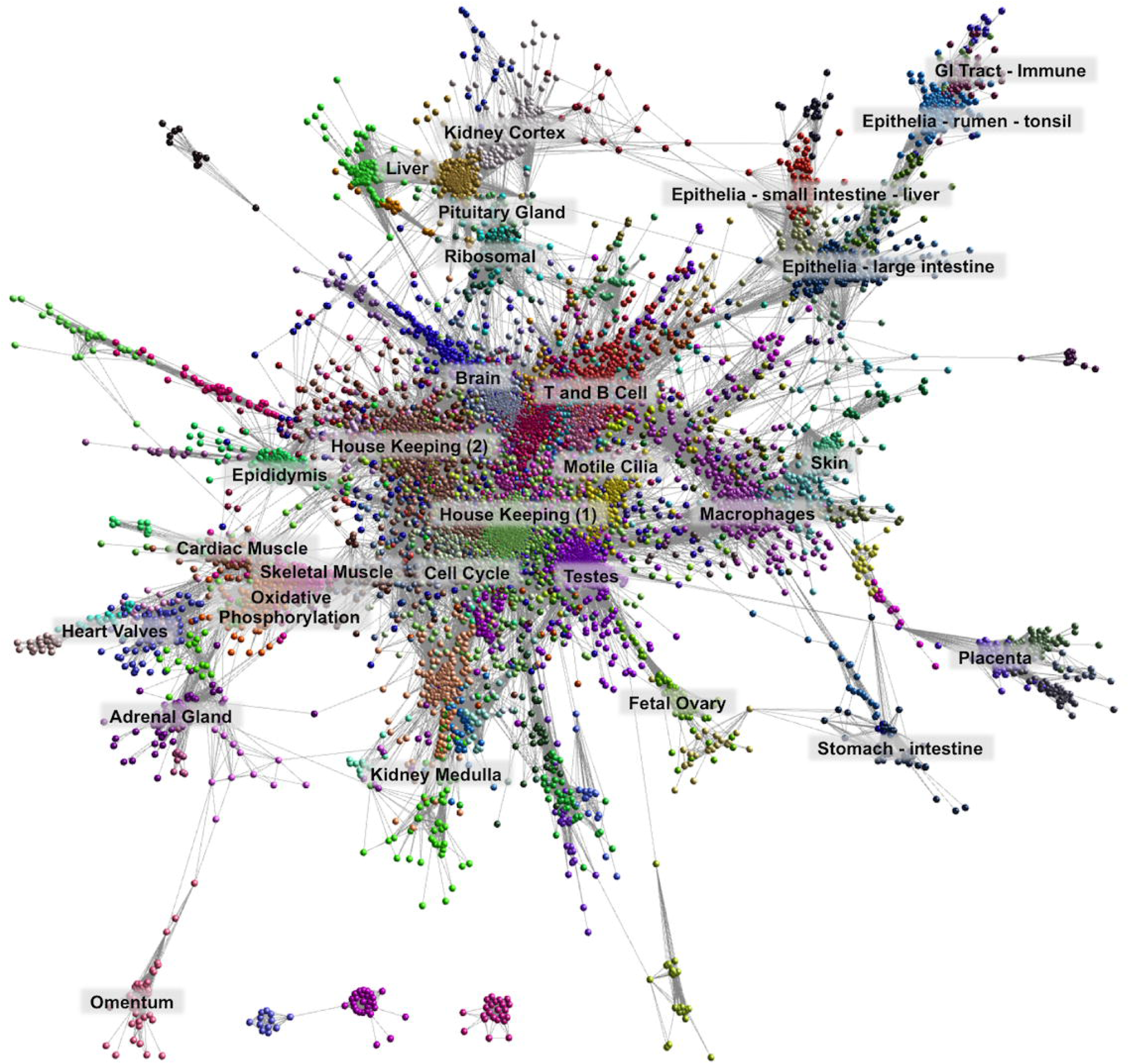
Network visualisation and clustering of the sheep gene expression atlas. A three-dimensional visualisation of a Pearson correlation gene-to-gene graph of expression levels derived from RNA-Seq data from analysis of sheep tissues and cells. Each node (sphere) in the graph represents a gene and the edges (lines) correspond to correlations between individual measurements above the defined threshold. The graph is comprised of 15,192 nodes (genes) and 811,213 edges (correlations ≥0.75). Co-expressed genes form highly connected complex clusters within the graph. Genes were assigned to groups according to their level of co-expression using the MCL algorithm.

**Fig 3:**
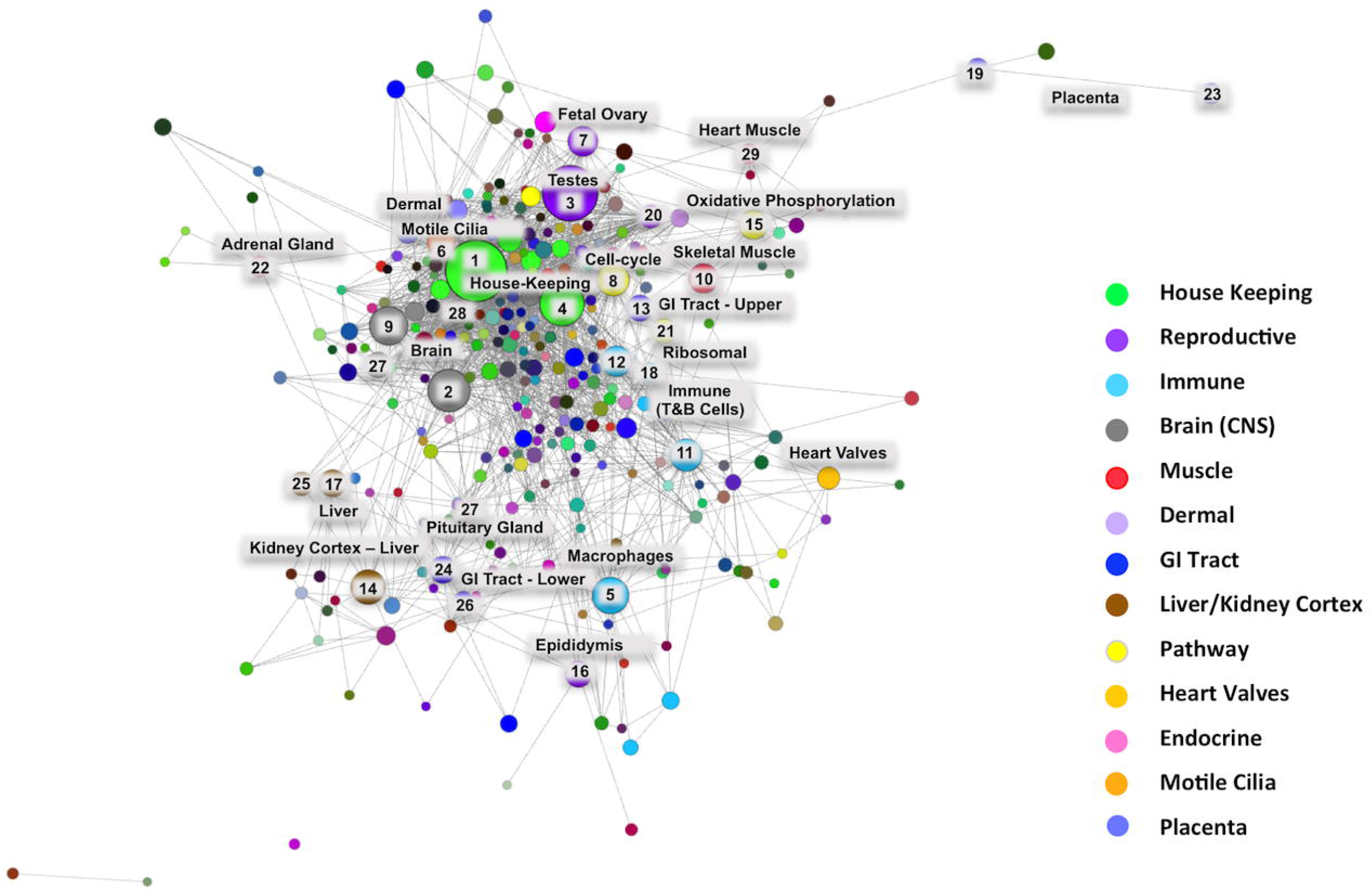
Collapsed node visualisation of the sheep gene expression atlas dataset in two-dimensions to illustrate the relative proportion of genes in each cluster. Includes 3104 nodes and 138,407 edges with a Pearson correlation value of *r*=0.75 and an MCL inflation (MCLi) value of 2.2. Nodes are coloured by tissue/cell type or for broader classes organ system. The largest clusters are numbered from 1 to 30 (see Table 3 for functional annotation). The largest clusters are dominated by either house-keeping genes (1 & 4) or genes associated with transcriptionally rich tissues or cell types, such as brain (2), testes (3) and macrophages (5).

**Table 3:** Tissue/cell/pathway association of the largest 50 network clusters in the sheep gene expression atlas dataset.

| Cluster ID | Number of Transcripts | Profile Description | Class | Sub-Class |
| --- | --- | --- | --- | --- |
| 1 | 1199 | General | House Keeping | House Keeping (1) |
| 2 | 987 | Brain | CNS | CNS |
| 3 | 658 | Testes – Adult | Reproduction | Gamete Production |
| 4 | 585 | General | House Keeping | House Keeping (2) |
| 5 | 351 | Macrophages | Immune | Macrophages |
| 6 | 350 | Fallopian Tube > Testes | Cilia | Motile Cilia |
| 7 | 284 | Fetal Ovary > Adult Testes | Early Development | Reproduction |
| 8 | 276 | Many Tissues - Highly Variable | Pathway | Cell Cycle |
| 9 | 265 | Fetal Brain > Adult Brain | CNS | CNS |
| 10 | 247 | Skeletal Muscle > Oesophageal Muscle | Musculature | Skeletal Muscle |
| 11 | 219 | Lymph Nodes > Blood > Not Macrophages | Immune | T-Cell and B-Cell |
| 12 | 215 | Thymus > Salivary Gland | Immune | T-Cell |
| 13 | 186 | Fore-Stomachs > Tonsil > Skin | GI Tract | Ruminal Epithelium |
| 14 | 183 | Kidney Cortex > Liver | Renal | Kidney Cortex |
| 15 | 182 | General but not even - highest in muscle | Pathway | Oxidative Phosphorylation |
| 16 | 158 | Epididymis > Vas Deferens | Reproduction | Male |
| 17 | 153 | Liver | Liver | Liver (Hepatocytes) |
| 18 | 145 | Peyer's Patch, Ileum, Lymph Nodes, Blood | Immune | T-Cell and B-Cell |
| 19 | 134 | Placenta | Gestation | Placental Function |
| 20 | 119 | Epididymis > Testes > Vas Deferens | Reproduction | Male |
| 21 | 115 | General but not even | Pathway | Ribosomal |
| 22 | 102 | Adrenal Gland | Endocrine | Steroid Hormone Biosynthesis |
| 23 | 102 | Placenta | Gestation | Placental Function |
| 24 | 98 | Liver > Small Intestine | Liver | Liver (GI Tract) |
| 25 | 96 | Fetal Liver | Liver | Developing Liver |
| 26 | 92 | Small Intestine > Large Intestine | GI tract | GI Tract |
| 27 | 90 | Pituitary Gland | Endocrine | Hormone Synthesis |
| 28 | 85 | General highest in reproductive tissues and brain | Cilia | Primary Cilia |
| 29 | 77 | Heart | Musculature | Cardiac Muscle |
| 30 | 75 | Thyroid | Endocrine | Thyroxine Synthesis |
| 31 | 73 | Peyers Patch, Ileum, Lymph Nodes, Blood, Macrophages | Immune | T-Cell and B-Cell |
| 32 | 69 | Salivary Gland, Lymph Node, Blood, Small Intestine | Immune | T-Cell and B-Cell |
| 33 | 68 | Fore-Stomachs Adult - Not Neonates > AMs | GI Tract | Immune |
| 34 | 65 | Small Intestine > Large Intestine | GI tract | GI Tract |
| 35 | 61 | General - highest in brain | House Keeping | House Keeping (3) |
| 36 | 58 | Heart Valves | Cardiovascular | Extra Cellular Matrix |
| 37 | 58 | General - highest in blood | House Keeping | House Keeping (4) |
| 38 | 56 | General - highest in fetal brain | House Keeping | House Keeping (5) |
| 39 | 50 | GI Tract - highest in reticulum | GI Tract | GI Tract |
| 40 | 49 | General - highest in testes | House Keeping | House Keeping (6) |
| 41 | 49 | General - highest in ovary | House Keeping | House Keeping (7) |
| 42 | 47 | General - highest in brain | House Keeping | House Keeping (8) |
| 43 | 45 | Fore-Stomachs > Tonsil > Skin | GI Tract | Ruminal Epithelium |
| 44 | 45 | General - highest in testes | House Keeping | House Keeping (9) |
| 45 | 44 | Macrophage (BMDM + LPS) | Immune | Macrophages (LPS Response TNF) |
| 46 | 44 | Blood, Lymph Nodes | Immune | Blood |
| 47 | 42 | Large Intestine | GI Tract | GI Tract |
| 48 | 42 | General | Pathway | Histones |
| 49 | 41 | Reticulum and Rumen - very variable | GI Tract | Reticulum/Rumen |
| 50 | 36 | Adrenal Gland Medulla | Endocrine | Steroid Hormone Biosynthesis |

The majority of co-expression clusters included genes exhibiting a specific cell/tissue expression pattern (Fig 4A). There were a few exceptions, including the largest cluster (cluster 1), which contained ubiquitously expressed ‘house-keeping’ genes, encoding proteins that are functional in all cell types. The high proportion of unannotated genes (24% of the 1199 genes) in cluster 1 may reflect the focus of functional genomics on genes exhibiting tissue specific expression, and inferred function in differentiation, leaving those with a house-keeping function uncharacterised [40]. With a few exceptions, the remaining co-expression clusters were comprised of genes exhibiting either expression only in a distinct tissue or cell type expression pattern e.g. macrophages (cluster 5) (Fig 4B (i)) and fetal ovary (cluster 7) (Fig 4B (ii)), or a broader expression pattern associated with a cellular process e.g. oxidative phosphorylation (cluster 15) (Fig 4B (iii)). Some co-expression is shared between two or more organ systems, associated with known shared functions. For example, cluster 15, exhibiting high expression in liver and kidney cortex, is enriched for expression of genes relating to the oxidation-reduction process, transmembrane transport, monocarboxylic acid catabolic process and fatty acid oxidation (S13 Table). It includes numerous genes encoding enzymes involved in amino acid catabolism (e.g. *AGXT, AGXT2*, *ASPDH*, *ACY1*, *EHHADH*, *DPYD*, *DAO*, *DDO*, *HAO1*, and *HPD*) and the rate-limiting enzymes of gluconeogenesis (*PCK1*, *PC*, *ALDOB* and *G6PC*). The contributions of kidney and liver to amino acid turnover and gluconeogenesis are well known in humans [41] and rodents [42]. These observations suggest that the shared catabolic pathways of liver and kidney cortex are largely conserved in sheep, but detailed curation of the genes in this cluster could provide further specific insights. Alanine aminotransferase (*ALT1* synonym *GPT1*), which generates alanine from the breakdown of amino acids in muscle and is transported to the liver for gluconeogenesis, is highly-expressed in muscle as expected. The glutaminase genes, required for the turnover of glutamine, are absent from tissue or cell type specific clusters; the liver-specific enzyme *GLS2* is also expressed in neuronal tissues, as it is in humans [43, 44].

**Fig 4:**
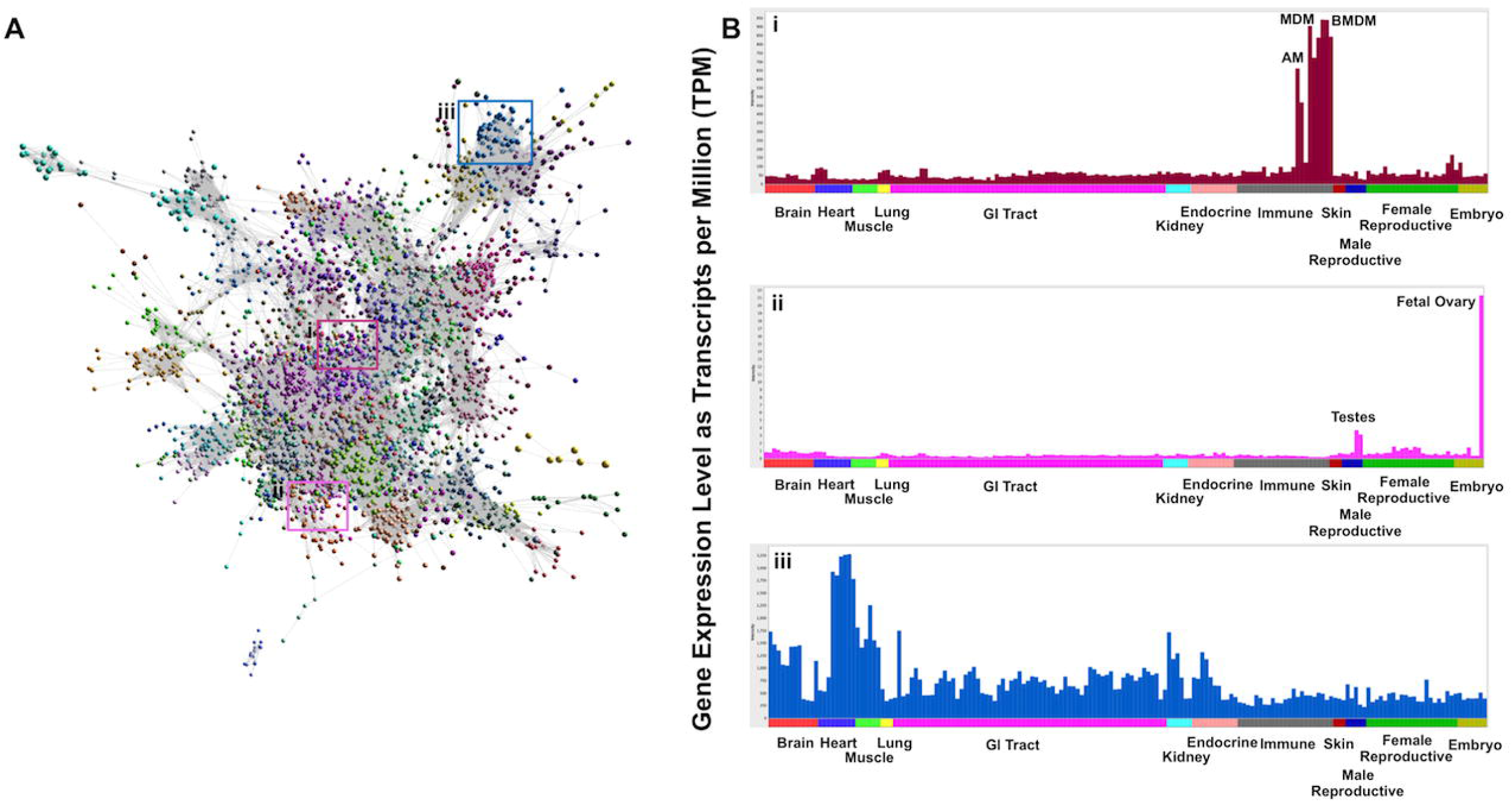
Interrogation of the underlying expression profiles allows regions of the graph to be associated with specific tissues or cell types. **A** A three-dimensional visualisation of a Pearson correlation gene to gene network graph (*r*=0.75, MCLi=2.2). Samples of each tissue and cell type from each breed and developmental stage are averaged across individuals for ease of visualisation. Histograms of the averaged expression profile (averaged across individuals for each tissue and cell type for ease of visualisation) of genes in selected clusters are given on the right: **B (i)** profile of cluster 5 genes whose expression is highest in macrophages; **(ii)** profile of cluster 7 genes whose expression is highest in fetal ovary and testes; **(iii)** or a broader expression pattern associated with a cellular process e.g. oxidative phosphorylation (cluster 15). Note that there may be a degree of variation in the expression pattern of individual genes within a cluster which is masked when average profiles are displayed.

The tissue-specific expression patterns observed across clusters showed a high degree of similarity to those observed for pig [6], human and mouse [8, 9]. In some cases we were able to add functional detail to clusters of genes previously observed in pig and human. For example, genes within cluster 6 showed high expression in the fallopian tube and to a lesser extent the testes. Significantly enriched GO terms for cluster 6 included cilium (p=4. 1×10^-8^), microtubule motor activity (p=2.9×10^-11^) and motile cilium (p=2.8×10^-19^) suggesting the genes expressed in cluster 6 are likely to have a function related to motile cilia in sperm cells and the fallopian tube. Cluster 6 in the sheep atlas dataset corresponds to cluster 9 in the pig gene expression atlas [6]. Similarly, significantly enriched GO terms for genes in cluster 28 included primary cilium (p=7.5×10^-21^) and ciliary basal body (p=1.9×10^-12^), indicating the genes in this cluster were associated with the function of primary cilia. Genes within this cluster showed a relatively wide expression pattern, with brain and reproductive tissues exhibiting the highest expression. For both clusters associated with cilial function, significantly enriched GO terms also included cell-cycle related cellular processes and cell-cycle associated genes, supporting the link between cilia and the cell-cycle [45-47].

### Cellular Processes

The genes within some clusters, rather than being linked to the function of a particular tissue or cell type, showed varying levels of expression across multiple tissues, suggesting their involvement in a universal cellular process (pathway). Significantly enriched GO terms for genes in cluster 8, for example, included ‘cell cycle checkpoint’ (p=1x10^-17^) and ‘mitotic cell cycle’ (p=<1x10^-30^). The variation in expression of these genes across tissues likely reflects variation in the proportion of mitotically active cells. In the same way, it is possible to extract a similar cluster from large cancer gene expression data sets, correlating with their proliferative index [48]. Expression of genes in cluster 15 (n=182) was detectable in most ovine tissues and cells but with strongly-enriched expression in skeletal and cardiac muscle. The pig gene expression atlas [6] highlighted an oxidative phosphorylation cluster and a mitochondrial/tricarboxylic acid (TCA) cluster. This functional grouping is merged in cluster 15. The majority of the transcripts in cluster 15 are present within the inventory of mitochondrial genes in humans and mice [49], but the reciprocal is not true since many other genes encoding proteins that locate to mitochondria were not found in cluster 15. Many mitochondrial proteins are unrelated to oxidative phosphorylation *per se*, and are enriched in other tissues including liver and kidney (including mitochondrial enzymes of amino acid and fatty acid catabolism, see above) and intestine.

Cluster 15 also contains several genes associated with myosin and the sarcoplasmic reticulum which may indicate some level of coordination of their function with the oxidative metabolism of glucose. The majority of genes in the corresponding cluster in pig [6] were also present in this cluster with a few notable additions including dihydrolipoamide dehydrogenase (*DLD*), which encodes a member of the class-I pyridine nucleotide-disulfide oxidoreductase family, and carnitine acetyltransferase (*CRAT*), a key enzyme in the metabolic pathway in mitochondria [50]. We were able to assign gene names to the following genes associated with oxidative phosphorylation complex I in pig: *NDUFA9* (ENSOARG00000009435), *NDUFB1* (ENSOARG00000020197), *NUDFB8* (assigned to ENSOARG00000015378), and *NDUFC2* (ENSOARG00000006694). The gene name *PTGES2* (prostaglandin E2 synthase 2) was assigned to ENSOARG00000010878, a gene that was associated with fatty acid (long-chain) beta oxidation in pig [6] but previously unannotated in sheep. Similarly, we assigned the gene name *PDHB* (pyruvate dehydrogenase E1 component subunit beta), a member of the pyruvate dehydrogenase complex also previously unannotated in sheep, to ENSOARG00000012222. The inclusion of the PDH complex, as well as the mitochondrial pyruvate carriers *MPC1* and *MPC2*, in the muscle-enriched cluster 15 reflects the fact that glucose, giving rise to pyruvate, is the preferred fuel for oxidative metabolism in muscle [51].

By using comparative clustering information in the pig and the “guilt by association” principle we were able to assign with confidence gene names and putative function to the majority of unannotated genes in clusters 8 (cell-cycle) and 15 (oxidative phosphorylation) (S13 Table). Expression of cell cycle and metabolic genes has recently been shown to be positively correlated with dry matter intake, ruminal short chain fatty acid concentrations and methane production in sheep [52]. In the same study a weak correlation between lipid/oxo-acid metabolism genes and methane yield was also identified suggesting that the newly unannotated genes in these clusters are likely to be relevant in addressing methane production in ruminants [52].

### The GI Tract

Stringent coexpression clustering requires that each transcript is quantified in a sufficiently large number of different states to establish a strong correlation with all other transcripts with which it shares coordinated transcription and, by implication, a shared function or pathway. The impact of this approach was evident from the pig gene expression atlas [6] which was effective at dissecting region-specific gene expression in the GI tract. We have generated a comparable dataset for sheep. In ruminants, the rumen, reticulum and omasum are covered exclusively with stratified squamous epithelium similar to that observed in the tonsil [18, 21]. Each of these organs has a very distinctive mucosal structure, which varies according to region sampled [53]. A network cluster analysis of regions of the GI tract from sheep has been published [21] using the Texel RNA-Seq dataset [18]. These co-expression clusters are broken up somewhat in this larger atlas, because many of the genes that are region-specific in the GI tract are also expressed elsewhere. We have, in addition, expanded the available dataset for the GI tract to include samples from neonatal and juvenile lambs.

The postnatal development of the sheep GI tract is of particular interest because of the pre-ruminant to ruminant transition, which occurs over 8 weeks from birth. Genes in cluster 33 showed low levels of expression in neonatal lambs and a gradual increase into adulthood. Enriched GO terms for this cluster include regulation of interleukin 6 (IL6) production (p=.0016) and keratinocyte differentiation (p=1.7×10^-8^) (S12 Table). The cluster includes genes such as *HMGCS2*, *HMGCL* and *BDH1*, required for ketogenesis, an essential function of rumen metabolism, as well as *CA1* (carbonic anhydrase 1), implicated in the rumen-specific uptake of short chain fatty acids. The cluster does not contain any of the solute carriers implicated in nutrient uptake in the rumen, suggesting that these are more widely-expressed and/or regulated from multiple promoters [21]. The only carrier that is rumen-enriched is *SLC9A3* (also known as *NHE3*), the key Na-H antiporter previously implicated in rumen sodium transport in both sheep and cattle [54]. Other genes in cluster 33, for example, *IL36A* and *IL36B*, for example, are thought to influence skin inflammatory response by acting on keratinocytes and macrophages and indirectly on T-lymphocytes to drive tissue infiltration, cell maturation and cell proliferation [55]. Many of these genes might also be part of the acute phase immune response, by regulating production of key cytokines such as *IL-6* and thus mediating activation of the NF-κB signaling pathways. Nuclear factor (NF)-κB and inhibitor of NF-κB kinase (IKK) proteins regulate innate- and adaptive-immune responses and inflammation (reviewed in [56]). Expression of many of these genes is likely to change as the immune system develops which we will describe in detail in a dedicated network cluster analysis of the GI tract developmental time series dataset. The genes in this cluster therefore appear to be involved both in the onset of rumination and in innate immunity (which could be associated with the population of the rumen microbiome).

### Innate and Acquired Immunity

Several clusters exhibited a strong immune signature. Clusters 11, 12, 18, 31 and 32, for example, contained genes with a strong T-lymphocyte signature [57] with high levels of expression in immune cell types and lymphoid tissues. Significantly enriched GO terms for cluster 12, for example, included T-cell differentiation (p=2.10x10^-11^), immune response (p=0.00182) and regulation of T-cell activation (p=0.00019) (S12 Table). Manual gene annotation using Ensembl IDs within this cluster revealed the majority were T-cell receptors and T-cell surface glycoproteins (S14 Table). Interestingly, ENSOARG00000008993 represents a gene with no orthologues to other species within the Ensembl database, but partial blast hits to T-lymphocyte surface antigen *Ly-9* in mouflon, goat, bison and buffalo in the NCBI database. The ‘true’ gene LY9, a member of the signalling lymphocyte activation molecule (SLAM) family [58], is also unannotated in sheep and is assigned to ENSOARG00000008981, having multiple orthologues in other placental mammals. We have assigned ENSOARG00000008993 the gene name ‘LY9-like’ and the symbol *LY9L*, and suggest this transcript plays a role in T-lymphocyte pathogen recognition.

Other immune clusters exhibited a macrophage-specific signature, with subsets highly expressed in alveolar macrophages (AMs), monocyte derived macrophages (MDMs) and bone marrow derived macrophages (BMDMs) (cluster 5) and two defined clusters of genes induced in BMDMs stimulated with LPS (cluster 45 and 52). Known macrophage-specific surface markers, receptors and proinflammatory cytokines predominated in these clusters, in addition to numerous unannotated genes, with as yet undefined but probable immune function (S15 Table). For example, the *CD63* antigen, which mediates signal transduction events, was assigned to ENSOARG00000011313 and *BST2* (bone marrow stromal cell antigen 2), which is part of the interferon (IFN) alpha/beta signaling pathway, to ENSOARG00000016787. A third cluster of LPS-inducible genes in macrophages, cluster 64, contained a subset of the IFN-inducible antiviral effector genes, including *DDX58*, *IFIT1*, *IFIT2*, *MX1*, *MX2*, *RSAD2* and *XAF1*, which are induced in mouse and humans through the MyD88-independent *TLR4* signaling pathway via autocrine *IFNB1* signaling (reviewed in [59]). Many other components of this pathway identified in LPS-stimulated human macrophages [60] were either not annotated, or not clustered, and will be the target of detailed annotation efforts in the macrophage dataset.

Significantly enriched GO terms for the macrophage-specific cluster 5 included ‘response to lipopolysaccharide’ (p=7.2×10^-7^), and ‘toll-like receptor signaling pathway’ (p=3.2×10^-5^). Many of the genes in this cluster are known components of the innate immune response in mammals. Interleukin 27 (*IL-27*), is a heterodimeric cytokine which has pro- and anti-inflammatory properties and a diverse effect on immune cells including the regulation of T-helper cell development, stimulation of cytotoxic T-cell activity and suppression of T-cell proliferation [61]. *ADGRE1* encodes the protein EGF-like module-containing mucin-like hormone receptor-like 1 (*EMR1* also known as F4/80), a classic macrophage marker in mice [62]. Several genes in cluster 5 encode proteins exclusively expressed in macrophages and monocytes. One such gene, *CD163*, encodes a member of the scavenger receptor cysteine-rich (SRCR) superfamily, which protects against oxidative damage by the clearance and endocytosis of hemoglobin/haptoglobin complexes by macrophages, and may also function as an innate immune sensor of bacteria [63].

One of the largest macrophage populations in the body occupies the lamina propria of the small and large intestine [64]. They are so numerous that the expression of macrophage-related genes can be detected within the total mRNA from intestine samples. As noted previously in the pig, one can infer from the expression profiles that certain genes that are highly-expressed in AMs are repressed in the intestinal wall [6]. We proposed that such genes, which included many c-type lectins and other receptors involved in bacterial recognition, were necessary for the elimination of inhaled pathogens, where such responses would be undesirable in the gut [6]. In the sheep, there was no large cohort of receptors that showed elevated expression in AMs relative to MDMs or BMDMs, and that were absent in the gut wall. Only a small cluster (115) of 13 genes showed that profile, including the phagocytic receptor *VSIG4* (CRiG), which is a known strong negative regulator of T-cell proliferation and *IL2* production [65] and *SCIMP*, recently identified as a novel adaptor of Toll-like receptor signaling that amplifies inflammatory cytokine production [66]. Six previously unannotated genes within this small cluster included the E3 ubiquitin ligase, *MARCH1*, and likely members of the paired immunoglobulin type receptor and SIGLEC families, which cannot be definitively assigned as orthologues.

Interestingly, macrophage colony-stimulating factor receptor (*CSF1R)*, which controls the survival, proliferation and differentiation of macrophage lineage cells [67, 68], was not within the macrophage-specific cluster 5. Instead, *CSF1R* was in a small cluster (cluster 102) along with several other macrophage-specific genes including the C1Q complex. As in humans and mice [9, 12, 69], *CSF1R* was also expressed in sheep placenta. In humans and mice, placental (trophoblast) expression is directed from a separate promoter [11]. The small number of genes co-expressed with *CSF1R* are likely either co-expressed by trophoblasts as well as macrophages (as is C1Q in humans, see BioGPS (http://biogps.org/dataset/GSE1133/geneatlas-u133a-gcrma/) [9, 70]), or highly-expressed in placenta-associated macrophages.

### Early Development and Reproduction

The sheep gene expression atlas dataset includes multiple libraries from early developmental time points. Three of the larger clusters of co-expressed genes showed high levels of expression in the fetal ovary (cluster 7), fetal brain (cluster 9) and fetal liver (cluster 25). ‘Testis-specific’ genes, particularly those involved in meiosis and gametogenesis, might also be expressed in the fetal ovary undergoing oogenesis [71, 72]. Our dataset from sheep appears to validate this hypothesis, since genes within cluster 7 exhibited higher levels of expression in the fetal ovary and to a lesser extent the testes. Several genes were expressed both in the testes and the fetal ovary including, testis and ovary specific PAZ domain containing 1 (*TOPAZ1*), which has been shown in sheep to be expressed in adult male testes and in females during fetal development with a peak during prophase I of meiosis [73]. Also, fetal and adult testis expressed 1 (*FATE1*), which is strongly expressed in spermatogonia, primary spermatocytes, and Sertoli cells in seminiferous tubules in mouse and humans [74]. Significantly enriched GO terms for genes within cluster 7 included ‘female gonad development’ (p=4.9×10^-6^), ‘spermatogenesis’ (p=4.6×10^-8^) and ‘growth factor activity’ (p=5×10^-5^) (S12 Table).

Several important genes for embryonic development were also co-expressed in cluster 7. The germ-cell specific gene SRY-box 30 (*SOX30*) encodes a member of the SOX (SRY-related HMG-box) family of transcription factors involved in the regulation of embryonic development and in the determination of cell fate [75]. Growth differentiation factor 3 (GDF3) encodes a protein required for normal ocular and skeletal development. Although it is a major stem cell marker gene [76], it has not previously been linked to germ cell expression. Similarly, POU class 5 homeobox 1 (*POU5F1*) encodes a transcription factor containing a POU homeodomain that controls embryonic development and stem cell pluripotency [76] but is also required for primordial germ cell survival [77]. The expression of these genes in tissues containing germ cells in sheep suggests they contribute to meiosis and cellular differentiation. These observations illustrate the utility of the sheep as a non-human model for the study of gametogenesis.

Cluster 7 also includes two related oocyte-derived members of the transforming growth factor-β (*TGFB1*) superfamily, growth differentiation factor 9 (*GDF9*) and bone morphogenetic protein 15 (*BMP15*), which are essential for ovarian follicular growth and have been shown to regulate ovulation rate and influence fecundity in sheep [78, 79]. Lambing rate is an important production trait in sheep and can vary between breeds based on single nucleotide polymorphism (SNP) mutations in key genes influencing ovulation rate (reviewed in [79, 80]). A number of the known fertility genes in sheep (reviewed in [81, 82]), such as the estrogen receptor (*ESR*) and the Lacaune gene (*B4GALNT2*) were not present in this cluster 7, which may be because they are not expressed in the ovary at the time points chosen for this study. Detailed analysis of the expression of key genes during early development in the fetal ovary in comparison with the ovary from the adult and gestating ewes may provide additional insights.

### Sex Specific Differences in Gene Expression

Sex-specific differences in gene expression have been reported in humans [83, 84] mice [85, 86], cattle [87, 88] and pigs [89, 90]. We examined male and female biased gene expression in the sheep atlas dataset by calculating the average TPM per sex for each gene and the female:male expression ratio (S3 Dataset). Twenty genes exhibited strongly sex biased expression (S16 Table); 13 were female-enriched and 7 were male-enriched. Among the male enriched genes was thyroid stimulating hormone beta (*TSHB*), which is expressed in thyrotroph cells in the pituitary gland and part of a neuro-endocrine signaling cascade in sheep [91]. Expression of *TSHB* in the pituitary gland of male BFxT was 3.6-fold higher than in female BFxT sheep. A similar sex bias has been observed in rats in which males exhibit significantly higher *TSHB* expression in the pituitary gland than females [92].

Other genes exhibiting similarly large sex specific fold-changes included keratin 36 (*KRT36*) which was expressed 6.6-fold higher in the reticulum of male relative to female sheep and *VSIG1* (V-Set and immunoglobulin domain containing 1), which is required for the differentiation of glandular gastric epithelia [93]. *VSIG1* showed 4-fold greater expression in the female pylorus relative to the male. An unannotated gene ENSOARG00000020792 exhibited large fold change in male biased expression in immune tissues including popliteal and prescapular lymph node, tonsil and Peyer’s patch. This gene has a detectable blast hit to “immunoglobulin kappa-4 light chain variable region” and is a 1:1 orthologue with an unannotated gene in cattle, ENSBTAG00000045514, with ≥70% reciprocal percentage identity and conservation of synteny. The dN/dS for ENSOARG00000020792 suggests it is evolving rapidly (dN/dS > 2). Male-biased genes are known to evolve quickly, as are immune genes [94]. GO term enrichment for the set of genes with five-fold sex-biased expression in at least one BFxT tissue (S17 Table) revealed that the genes enriched in females were predominately involved with the immune reponse while gene enriched in the male were broadly associated with muscle and connective tissue. This is likely to reflect inherent differences between the two sexes in allocation of resources towards growth or immunity. Genes exhibiting sex specific expression might therefore be relevant in sexual diamorphism in disease susceptibility for example, and warrant further investigation.

### Differential Expression of Genes between the Texel and BFxT

The majority of commercially-produced livestock are a cross of two or more different production breeds with distinct desired traits [95]. For example, in the UK, the crossing of lighter upland sheep breeds with heavier lowland meat breeds optimises carcass quality, lambing rate, growth rate and survivability [95]. In developing countries, sustainable crossing of indigenous small ruminants with elite western breeds is one approach to improve productivity [96, 97]. An RNA-Seq dataset of this size from an outbred cross of two disparate sheep breeds provides an opportunity to investigate differential gene expression in a purebred parental line and crossbred animals. We compared gene expression across tissues in the F_1_ crossbred (BFxT) animals (generated by crossing a Texel ram with a Scottish Blackface ewe; Fig 5A) with the purebred Texel animals included in the previous sheep gene expression atlas dataset [18].

**Fig 5:**
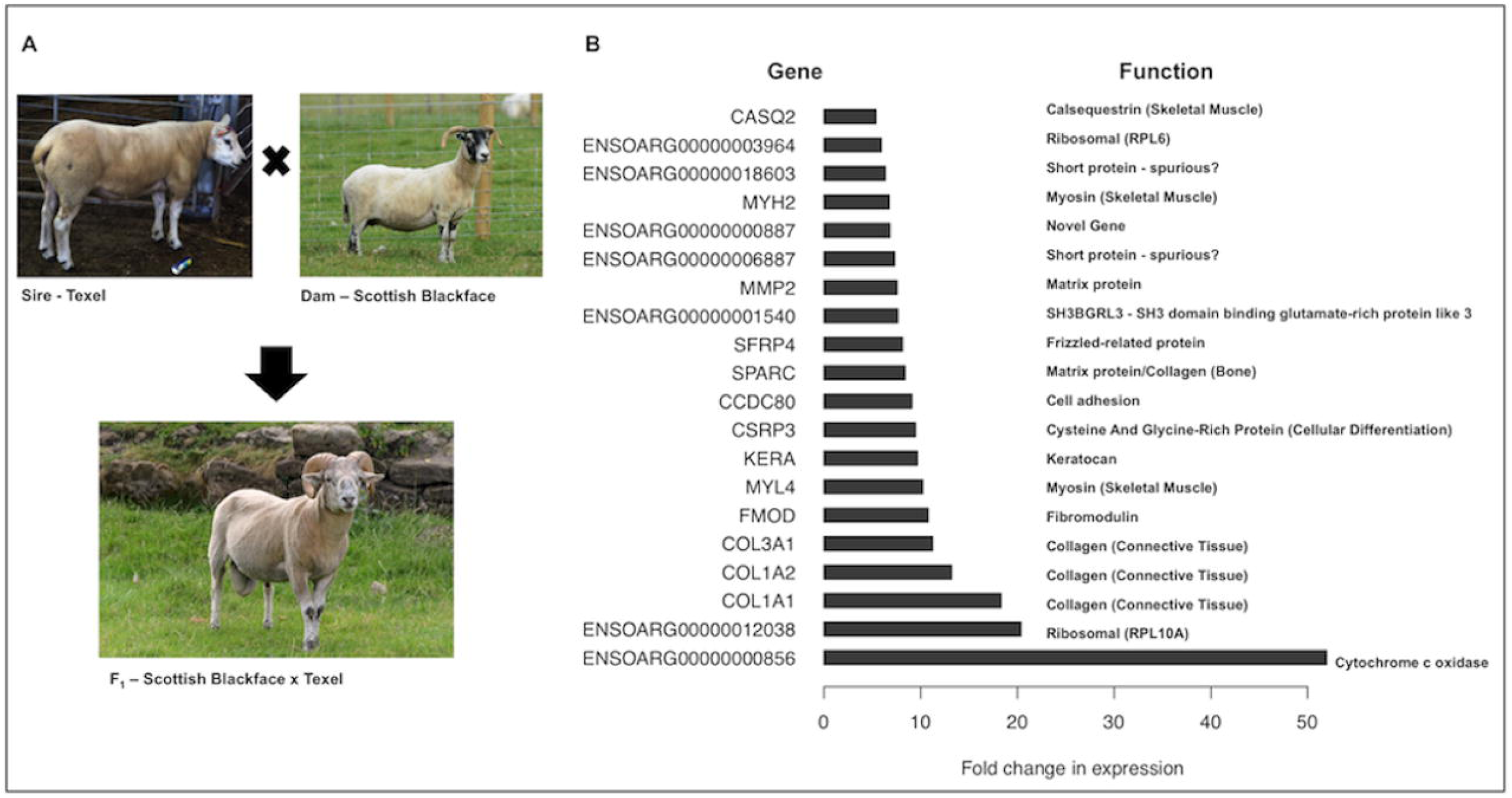
Upregulation of genes in the crossbred BFxT relative to the purebred Texel. **A** A Texel sire was crossed with Scottish Blackface dam to create the F1 Texel x Scottish Blackface individuals. **B** The top 20 genes showing the greatest up-regulation (as absolute fold change) between the crossbred BFxT and purebred Texel individuals. Genes associated with the function of skeletal muscle and connective tissue are indicated.

A gene was considered differentially expressed (DE) between the purebred Texel and hybrid BFxT if (a) it was expressed at ≥1 TPM in both Texel and BFxT (considering TPM to be the mean of all replicates per tissue), (b) the fold change (ratio of BFxT TPM to Texel TPM) was ≥2 in ≥25% of the tissues in which expression was detected (stipulating no minimum number of tissues, but noting that 23 tissues are common to Texel and BFxT), and (c) the fold change was ≥5 in at least 1 tissue. Fold changes of all genes expressed at ≥1 TPM in both breeds are given in S18 Table. The GO terms enriched in the set of DE genes (n=772) with higher expression in the BFxT than the Texel were predominantly related either to muscle or brain function (S19 Table). The top 20 genes showing the largest up-regulation (shown as absolute fold-change) in the BFxT relative to the purebred Texel sheep are illustrated in Fig 5B. Enriched molecular function GO terms for the set of genes differentially expressed between BFxT and Texel sheep include ‘iron ion binding’ (p=3.6×10^-4^), and ‘cytoskeletal protein binding’ (p=7.8×10^-6^), biological process terms include ‘cellular iron ion homeostasis’ (p=6.3×10^-4^) and cellular component terms include ‘sarcomere’ (p=6.6×10^-7^) and ‘collagen trimer’ (p=5.5×10^-7^) (S19 Table).

Numerous genes with structural, motor and regulatory functions were highly expressed in BFxT compared to Texel bicep muscle, with approximately 5- to 18-fold expression increases for various members of the collagen (*COL1A1*, *COL1A2*, *COL3A1*) and myosin families (*MYH2*, *MYL4*), along with *CSRP3* (a mechanosensor) [98], *FMOD* (fibromodulin, a regulator of fibrillogenesis) [99], keratocan (*KERA*, a proteoglycan involved in myoblast differentiation) [100], matrix metalloproteinase 2 (*MMP2*, a proteolytic enzyme associated with muscle regeneration) [101], and calsequestrin 2 (*CASQ2*, one of the most abundant Ca^2+^-binding proteins in the sarcoplasmic reticulum, essential for muscle contraction) [102].

Genes enriched in muscle are of particular biological and commercial interest because Texel sheep exhibit enhanced muscling and less fat [103], due to a single nucleotide polymorphism (SNP) in the 3’ untranslated region of the myostatin gene *MSTN* (synonym *GDF-8*) which generates an illegitimate miRNA binding site resulting in translational inhibition of myostatin synthesis and contributing to muscular hypertrophy [24, 104]. Because heterozygotes have an intermediate phenotype, cross breeding of homozygous mutant Texel sheep with animals homozygous for the normal allele transmits something of the Texel muscle phenotype to the offspring. An effect on muscle synthesis in the BFxT animals could be related to the myostatin genotype; genes with higher expression in the Texel than in the cross may be targets for myostatin inhibition [105], while those with lower expression in the Texel than in the cross may be directly or indirectly activated by myostatin and hence involved in the cessation of muscle differentiation. In cattle the myostatin mutation is associated with the downregulation of collagen genes including *COL1A1* and *COL1A2* [105]. This is consistent with the observation that these genes have higher expression in the heterozygous BFxT animals than the Texel animals. Since myostatin also regulates muscle fibre type [106] by suppressing the formation of fast-twitch fibres, individuals homozygous for inactivating myostatin mustations are likely to exhibit increased fast-twitch fibres [107]. Many of the genes up-regulated in the BFxT relative to the Texel animals (e.g. *CSRP3* and *CASQ2*) are known to be specifically expressed in slow-twitch muscle [106, 108], and several down-regulated genes are associated with fast-twitch muscle (e.g. *TNNC2*, *TNNI2* and *SERCA1*) [109, 110]. Consequently, the difference between the cross-breed and pure Texel is in part attributable to an increased contribution of slow-twitch fibres, which in turn has been associated with desirable meat quality traits [111] highlighting the potential advantages of cross-breeding.

Enriched GO terms related to brain function include the ‘myelin sheath’ (p=6.1x10^-8^) and the ‘internode region of the axon’ (p=5.2x10^-5^) (S19 Table). Candidate genes of particular interest were expressed in the cerebellum (S18 Table). For instance, in the BFxT relative to the Texel animals, there were approximately 8-fold expression increases in cochlin (*COCH*, which regulates intraocular pressure) [112] and brevican (*BCAN*, functions throughout brain development in both cell-cell and cell-matrix interactions) [113, 114] and a 10-fold expression increase for myelin-associated oligodendrocyte basic protein, MOBP, which was previously unannotated in sheep (ENSOARG00000002491) and has a function in late-stage myelin sheath compaction [115]. A 15-fold expression increase was observed for oligodendrocytic paranodal loop protein (*OPALIN*, a transmembrane protein unique to the myelin sheath [116]), in the BFxT relative to the Texel and a 10-fold increase for another unannotated gene myelin basic protein (*MBP*), which we have assigned to ENSOARG00000004374 [117]. Although these examples of a neuroendocrine-specific effect of cross-breeding are speculative, they are of interest as Scottish Blackface sheep exhibit both improved neonatal behavioural development [118] and more extensive foraging behaviour than lowland breeds such as the Texel, travelling further distances, covering greater areas and exploring higher altitudes [119].

### Visualisation of the Expression Atlas

We have provided the BFxT sheep gene expression atlas as a searchable database in the gene annotation portal BioGPS (http://biogps.org/dataset/BDS_00015/sheep-atlas/). By searching the dataset via the following link (http://biogps.org/sheepatlas/) the expression profile of any given gene can be viewed across tissues. An example profile of the *MSTN* (*GDF-8*) myostatin gene from sheep is included in Fig 6. BioGPS allows comparison of expression profiles across species and links to gene information for each gene [70, 120, 121]. The Sheep Atlas BioGPS expression profiles are based on TPM estimates from the alignment-free Kallisto output for the BFxT libraries, averaged across samples from each developmental stage for ease of visualisation. It is important to note that there may be a degree of variation in the expression pattern of individual genes between individuals which is masked when the average profiles are displayed. In addition, to allow comparison between species BioGPS requires each gene have an Entrez ID, which is not the case for all genes in Oar v3.1 and as a consequence these genes do not have visualisable expression profiles in BioGPS. The expression profiles of the genes without Entrez IDs can be found in S1 Dataset and S2 Dataset.

**Fig 6:**
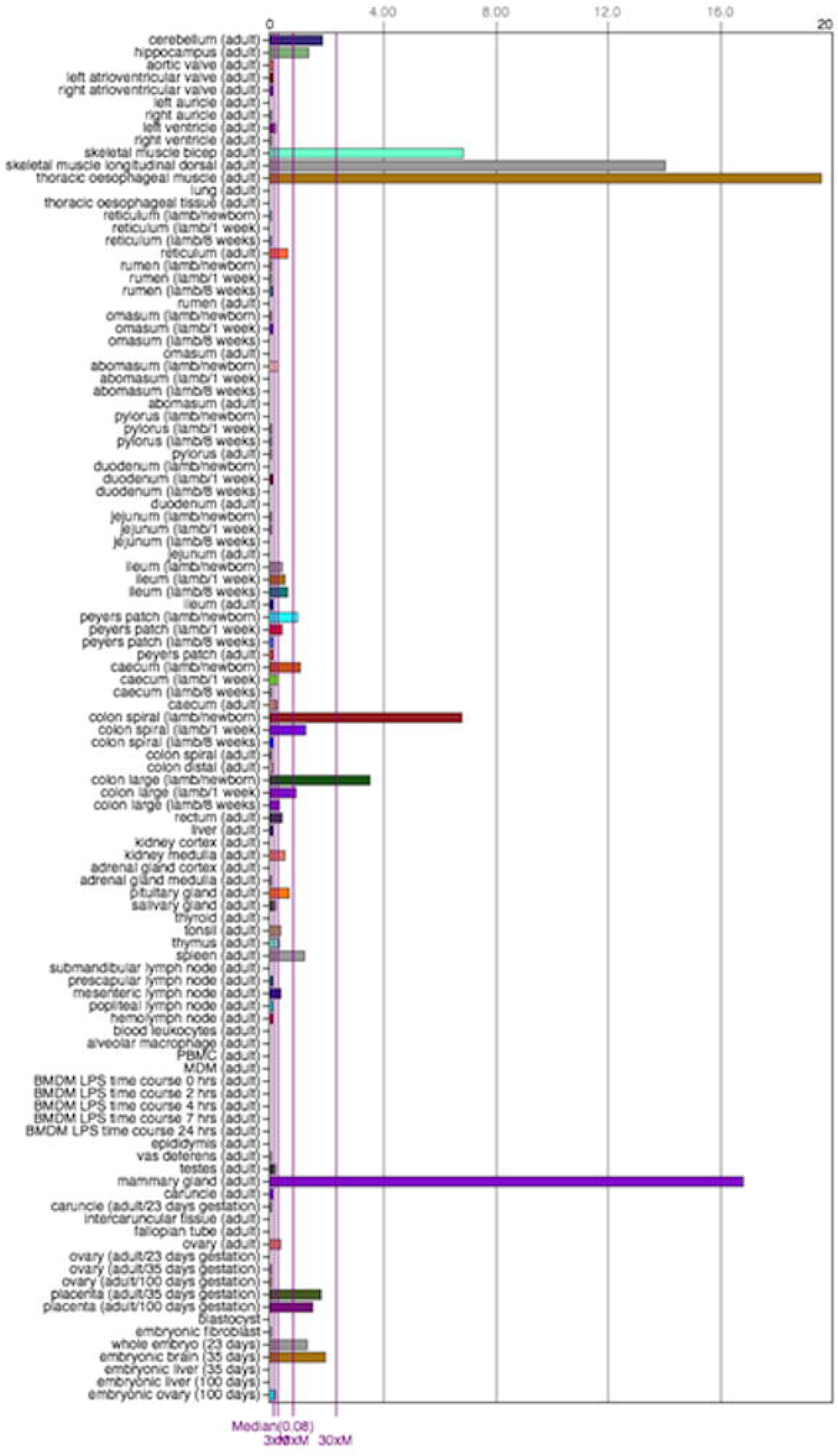
Screenshot of the representation of the profile of the sheep *GDF-8* (*MSTN*) gene within the BioGPS online portal. All data from the BFxT sheep gene expression atlas dataset are available through the BioGPS database (http://biogps.org/dataset/BDS_00015/sheep-atlas/). This provides a searchable database of genes, with expression profiles across tissues and cells for each gene displayed as histograms via the following link, http://biogps.org/sheepatlas/. Samples are coloured according to organ system, for example, Immune, GI Tract etc. The BioGPS platform supports searching for genes with a similar profile, allows access to the raw data, and links to external resources. It also provides the potential for comparative analysis across species, for example with the expression profiles for pig.

In parallel to the alignment-free Kallisto method, we also used an alignment-based approach to RNA-Seq processing, with the HISAT aligner [122] and StringTie assembler [123] (detailed in S1 Methods). These alignments will be published as tracks in the Ensembl genome browser in the short term and integrated into the next Ensembl genome release for sheep.

## Conclusions

This work describes the transcriptional landscape of the sheep across all major organs and multiple developmental stages, providing the largest gene expression dataset from a livestock species to date. The diversity of samples included in the sheep transcriptional atlas is the greatest from any mammalian species, including humans. Livestock provide an attractive alternative to rodents as models for human conditions in that they are more human-like in their size, physiology and transcriptional regulation, as well as being economically important in their own right. Non-human models are required to study the fundamental biology of healthy adult mammals and as such this dataset represents a considerable new resource for understanding the biology of mammalian tissues and cells.

In this sheep transcriptional atlas gene expression was quantified at the gene level across a comprehensive set of tissues and cell-types, providing a starting point for assigning function based on cellular expression patterns. We have provided functional annotation for hundreds of genes that previously had no meaningful gene name using co-expression patterns across tissues and cells. Future analysis of this dataset will use the co-expression clusters to link gene expression to observable phenotypes by highlighting the expression patterns of candidate genes associated with specific traits from classical genetic linkage studies or genome-wide association studies (GWAS). Gene expression datasets have been used in this way to characterise cell populations in mouse [22, 23] and in the biological discovery of candidate genes for key traits in sheep [124-126] and pigs [127, 128]. We have already utilised the dataset to examine the expression patterns of a set of candidate genes linked to mastitis resistance [129] in sheep, including comparative anlaysis with a recently available RNA-Seq dataset from sheep lactating mammary gland and milk samples [130]. The research community will now be able to use the sheep gene expression atlas dataset to examine the expression patterns of their genes or systems of interest, to answer many of the outstanding questions in ruminant biology, health, welfare and production.

Improving the functional annotation of livestock genomes is critical for biological discovery and in linking genotype to phenotype. The Functional Annotation of Animal Genomes Consortium (FAANG) aims to establish a data-sharing and research infrastructure capable of analysing efficiently genome wide functional data for animal species [26, 27]. This analysis is undertaken on a large scale, including partner institutions from across the globe, to further our understanding of how variation in gene sequences and functional components shapes phenotypic diversity. Analysis of these data will improve our understanding of the link between genotype and phenotype, contribute to biological discovery of genes underlying complex traits and allow the development and exploitation of improved models for predicting complex phenotypes from sequence information. The sheep expression atlas is a major asset to genome assembly and functional annotation and provide the framework for interpretation of the relationship between genotype and phenotype in ruminants.

## Methods

### Animals

Approval was obtained from The Roslin Institute’s and the University of Edinburgh’s Protocols and Ethics Committees. All animal work was carried out under the regulations of the Animals (Scientific Procedures) Act 1986. Three male and three female Scottish Blackface x Texel sheep of approximately two years of age were acquired locally and housed indoors for a 7-10 day “settling-in period” prior to being euthanased (electrocution and exsanguination). Nine Scottish Blackface x Texel lambs were born at Dryden Farm Large Animal Unit. Three neonatal lambs were observed at parturition and euthanised immediately prior to their first feed, three lambs were euthanised at one week of age prior to rumination (no grass was present in their GI tract) and three at 8 weeks of age once rumination was established. The lambs were euthanised by schedule one cranial bolt. To obtain developmental tissues six Scottish Blackface ewes were mated to a Texel ram and scanned to ensure successful pregnancy at 21 days. Two were euthanised at 23 days, two at 35 days and two at 100 days gestation (electrocution followed by exsanguination). Corresponding time points (day 23, 35 and 100) from gestating BFxT ewes mated with a Texel ram and euthanised as for the Blackface ewes were also included. All the animals, were fed *ad libitum* on a diet of hay and 16% sheep concentrate nut, with the exception of the lambs pre-weaning who suckled milk from their mothers. Details of the animals sampled are included in S1 Table.

### Tissue Collection

Tissues (95 tissues/female and 93 tissues/male) and 5 cell types were collected from three male and three female adult Scottish Blackface x Texel (BFxT) sheep at two years of age. The same tissues were collected from nine lambs, 3 at birth, 3 at one week and 3 at 8 weeks of age. Three embryonic time points were also included: three day-23 BFxT whole embryos, three BFxT day 35 embryos from which tissue was collected from each region of the basic body plan and three day 100 BFxT embryos from which 80 tissues were collected. Reproductive tissue from the corresponding time points from 6 BFxT ewes, 2 at each gestational time point, was also collected. In addition, 3 pools of 8 day 7 blastocysts from abattoir derived oocytes (of unknown breed) fertilized with Texel semen were created using IVF.

The majority of tissue samples were collected into RNAlater (AM7021; Thermo Fisher Scientific, Waltham, USA) and a subset were snap frozen, including lipid rich tissues such as adipose and brain. To maintain RNA integrity all tissue samples were harvested within an hour from the time of death. A detailed list of the tissues collected and sequenced can be found in S2 Table. Within the scope of the project we could not generate sequence data from all the samples collected and have archived the remainder for future analysis. Sample metadata, conforming to the FAANG Consortium Metadata standards, for all the samples collected for the sheep gene expression atlas project has been deposited in the BioSamples database under project identifier GSB-718 (https://www.ebi.ac.uk/biosamples/groups/SAMEG317052).

### Isolation of Cell Types

All cell types were isolated on the day of euthanasia. Bone marrow cells were isolated from 10 posterior ribs as detailed for pig [131]. BMDMs were obtained by culturing bone marrow cells for 7 days in complete medium: RPMI 1640, Glutamax supplement (35050-61; Invitrogen, Paisley, U.K.), 20% sheep serum (S2263; Sigma Aldrich, Dorset, U.K), penicillin/streptomycin (15140; Invitrogen) and in the presence of recombinant human CSF-1 (rhCSF-1: 10^4^ U/ml; a gift of Chiron, Emeryville, CA) on 100-mm^2^ sterile petri dishes, essentially as described previously for pig [131]. For LPS stimulation the resulting macrophages were detached by vigorous washing with medium using a syringe and 18-g needle, washed, counted, and seeded in tissue culture plates at 10^6^ cells/ml in CSF-1–containing medium. The cells were treated with LPS from *Salmonella enterica* serotype minnesota Re 595 (L9764; Sigma-Aldrich) at a final concentration of 100 ng/ml as previously described in pig [131] and harvested into TRIzol^®^ (Thermo Fisher Scientific) at 0, 2, 4, 7 and 24 h post LPS treatment before storing at -80°C for downstream RNA extraction.

PBMCs were isolated as described for pig [132]. MDMs were obtained by culturing PBMCs for 7 days in CSF-1–containing medium, as described above for BMDMs, and harvesting into TRIzol^®^ (Thermo Fisher Scientific). Alveolar macrophages were obtained by broncho-alveolar lavage of the excised lungs with 500ml sterile PBS (Mg^2+^ Ca^2+^ free) (P5493; Sigma Aldrich). The cells were kept on ice until processing. To remove surfactant and debris cells were filtered through 100uM cell strainers and centrifuged at 400 × *g* for 10 min. The supernatant was removed and 5ml red blood cell lysis buffer (420301; BioLegend, San Diego, USA) added to the pellet for 5 min; then the cells were washed in PBS (Mg^2+^ Ca^2+^ free) (P5493; Sigma Aldrich) and centrifuged at 400 × *g* for 10 min. The pellet was collected, resuspended in sterile PBS (Mg^2+^ Ca^2+^ free) (Sigma Aldrich), and counted. Alveolar macrophages were seeded in 6-well tissue culture plates in 2ml complete medium: RPMI 1640, Glutamax supplement (35050-61; Invitrogen), 20% sheep serum (S2263; Sigma Aldrich), penicillin/streptomycin (15140; Invitrogen) in the presence of rhCSF1 (10^4^ U/ml) overnight.

Blood leukocytes were isolated as described in [133]. Whole blood was spun at 500 x *g* for 10 min (no brake) to separate the buffy coats. These were then lysed in ammonium chloride lysis buffer (150mM NH_4_Cl, 10mM NaHCO_3_, 0.1mM EDTA) for 10 min on a shaking platform, then centrifuged at 4°C for 5 min at 500 x *g*. The resultant blood leukocyte pellets were stored in 1ml of RNAlater (Thermo Fisher Scientific) at -80°C.

To isolate embryonic fibroblasts we harvested a day 35 embryo whole and transferred to outgrowth media (DMEM, high glucose, glutamine, pyruvate (Thermo Fisher Scientific; 11995065), FBS (Fetal Bovine Serum) (Thermo Fisher Scientifc; 10500056), MEM NEAA (Thermo Fisher Scientific; 11140068), penicillin/streptomycin (Invitrogen), Fungizone (Amphotericin B; Thermo Fisher Scientific; 15290018), Gentamicin (Thermo Fisher Scientific; 15750037)). In a sterile flow hood the head was removed and the body cavity eviscerated. The remaining tissue was washed 3 times in PBS (Mg^2+^ Ca^2+^ free) (Sigma Aldrich) with penicillin/streptomycin (Invitrogen). 5ml of Trypsin-EDTA solution (T4049; Sigma Aldrich) was added and the sample incubated at 37°C for 5 min then vortexed and incubated for an additional 5min at 37°C. 3ml of solution was removed and filtered through a 100uM cell strainer, 5ml of outgrowth media was then passed through the strainer and combined with the sample. The sample was centrifuged at 200 x *g* for 3 min and the pellet resuspended in 9ml of out growth media before splitting the sample between 3x T75 flasks. The process then was repeated for remaining 2ml of sample left after the digestion with Trypsin (above). Embryonic fibroblasts were incubated for 5-7 days (until 80-90% confluent) then harvested into TRIzol^®^ Reagent (Thermo Fisher Scientific).

### RNA Extraction and Library Preparation

RNA was extracted using the same method as the Roslin RNA-Seq samples included in the sheep genome project detailed in [18]. For each RNA extraction <100mg of tissue was processed. Care was taken to ensure snap frozen samples remained frozen prior to homogenisation, and any cutting down to the appropriate size was carried out over dry ice. Tissue samples were first homogenised in 1ml of TRIzol^®^ reagent (15596018; Thermo Fisher Scientific) with either CK14 (432-3751; VWR, Radnor, USA) or CKMIX (431-0170; VWR) tissue homogenising ceramic beads on a Precellys^®^ Tissue Homogeniser (Bertin Instruments; Montigny-le-Bretonneux, France). Homogenisation conditions were optimised for tissue type but most frequently 5000 rpm for 20 sec. Cell samples which had previously been collected in TRIzol^®^ Reagent (15596018; Thermo Fisher Scientific), were mixed by pipetting to homogenise. Homogenised (cell/tissue) samples were then incubated at room temperature for 5 min to allow complete dissociation of the nucleoprotein complex, 200μl BCP (1-bromo-3- chloropropane) (B9673; Sigma Aldrich) was added, then the sample was shaken vigorously for 15 sec and incubated at room temperature for 3 min. The sample was centrifuged for 15 min at 12,000 x *g*, at 4°C to separate the homogentate into a clear upper aqueous layer (containing RNA), an interphase and red lower organic layers (containing the DNA and proteins), for three min. DNA and trace phenol was removed using the RNeasy Mini Kit (74106; Qiagen Hilden, Germany) column purification, following the manufacturers instructions (Protocol: Purification of Total RNA from Animal Tissues, from step 5 onwards). RNA quantity was measured using a Qubit RNA BR Assay kit (Q10210; Thermo Fisher Scientific) and RNA integrity estimated on an Agilent 2200 Tapestation System (Agilent Genomics, Santa Clara, USA) using the RNA Screentape (5067-5576; Agilent Genomics) to ensure RNA quality was of RIN^e^ > 7.

RNA-Seq libraries were prepared by Edinburgh Genomics (Edinburgh Genomics, Edinburgh, UK) and run on the Illumina HiSeq 2500 sequencing platform (Illumina, San Diego, USA). Details of the libraries generated can be found in S2 Table. A subset of 10 tissue samples and BMDMs at 0 h and 7h (+/-LPS) (Table 1), from each individual, were sequenced at a depth of >100 million strand-specific 125bp paired-end reads per sample using the standard Illumina TruSeq total RNA library preparation protocol (Ilumina; Part: 15031048, Revision E). These samples were chosen to include the majority of transcriptional output, as in [134]). An additional 40 samples from the tissues and cell types collected per individual (44/female and 42/male), were selected and sequenced at a depth of >25 million strand-specific paired-end reads per sample using the standard Illumina TruSeq mRNA library preparation protocol (poly-A selected) (Ilumina; Part: 15031047 Revision E).

In addition to the samples from the 6 adults, tissue was also collected from other developmental time points. The GI tract tissues collected from the 9 BFxT lambs, 3 at birth, 3 at one week of age and 3 at 8 weeks of age were sequenced at a depth of >25 million reads per sample using the Illumina mRNA TruSeq library preparation protocol (poly-A selected) as above. Of the early developmental time points, the three 23 day old embryos from BFxT sheep were sequenced at >100 million reads using the Illumina total RNA TruSeq library preparation protocol (as above), while the other embryonic samples and the ovary and placenta from the gestating ewes were sequenced at a depth of >25 million reads per sample using the Illumina mRNA TruSeq library preparation protocol (as above). In addition, three libraries were generated using the NuGen Ovation Single Cell RNA-Seq System (0342-32-NUG; NuGen, San Carlos, USA) from pooled samples of 8 blastocysts (as in [135]), and sequenced at a depth of >60 million reads per sample. A detailed list of prepared libraries, including library type can be found in S2 Table.

To identify spurious samples we used sample-to-sample correlation, of the transposed data from S1 Dataset, in Miru (Kajeka Ltd) [37]. The sample-to-sample graph is presented in S2 Fig. The expression profiles of any samples clustering unexpectedly (i.e. were not found within clusters of samples of the same type/biological replicate) were examined in detail. Generally the correlation between samples was high, although 4 spurious samples and 4 sets of swapped samples were identified. These samples were either relabeled or removed as appropriate.

### Data Quality Control and Processing

Raw data is deposited in the European Nucleotide Archive under study accession number PRJEB19199 (http://www.ebi.ac.uk/ena/data/view/PRJEB19199). The RNA-Seq data processing methodology and pipelines are described in detail in S1 Methods. For each tissue a set of expression estimates, as transcripts per million (TPM), were obtained using the high speed transcript quantification tool Kallisto [29]. In total, the expression atlas utilised approximately 26 billion (pseudo) alignments (S3 Table), capturing a large proportion of protein-coding genes per tissue (S20 Table).

The accuracy of Kallisto is dependent on a high quality index (reference transcriptome) [29], so in order to ensure an accurate set of gene expression estimates we employed a ‘two-pass’ approach. We first ran Kallisto on all samples using as its index the Oar v3.1 reference transcriptome. We then parsed the resulting data to revise this index. This was in order to include, in the second index, those transcripts that should have been there but were not (i.e. where the reference annotation is incomplete), and to remove those transcripts that should not be there but were (i.e. where the reference annotation is poor quality and a spurious model has been introduced). For the first criterion, we obtained the subset of reads that Kallisto could not align, assembled those *de novo* into putative transcripts (S1 Methods), then retained each transcript only if it could be robustly annotated (by, for instance, encoding a protein similar to one of known function) and showed coding potential (S21 Table). For the second criterion, we identified those members of the reference transcriptome for which no evidence of expression could be found in any of the hundreds of samples comprising the atlas. These were then discarded from the index. Finally, this revised index was used for a second iteration of Kallisto, generating higher-confidence expression level estimates. This improved the capture rate of protein-coding genes (S22 Table). A detailed description of this process can be found in S1 Methods.

We complemented this alignment-free method with a conventional alignment-based approach to RNA-Seq processing, using the HISAT aligner [122] and StringTie assembler [123]. A detailed description of this pipeline is included in S1 Methods. This assembly is highly accurate with respect to the existing (Oar v3.1) annotation, precisely reconstructing almost all exon (96%), transcript (98%) and gene (99%) models (S23 Table). Although this validates the set of transcripts used to generate the Kallisto index, we did not use HISAT/StringTie to quantify expression. This is because a standardised RNA space is necessary to compare data from mRNA-Seq and total RNA-Seq libraries [31], which cannot be applied if expression is quantified via genomic alignment. Unlike alignment-free methods, however, HISAT/StringTie can be used to identify novel transcript models (S24 Table), particularly for ncRNAs, which will be described in detail in a dedicated analysis. We will publish the alignments from HISAT and Stringtie as tracks in the Ensembl genome browser in the short term and integrate the alignments into Ensembl and Biomart in the next Ensembl release for sheep.

### Inclusion of Additional RNA-Seq Datasets from Sheep

Additional RNA-Seq data was obtained from a previous characterisation of the transcriptome of 3 Texel sheep included in the release of the current sheep genome Oar v3.1 [18]. The dataset included tissues from an adult Texel ram (n=29), an adult Texel ewe (n=25) and their female (8-9 month old) lamb (n=28), plus a whole embryo (day 15 gestation) from the same ram-ewe pairing. The raw read data from the 83 Texel samples incorporated into this dataset and previously published in [18] is located in the European-Nucleotide Archive (ENA) study accession PRJEB6169 (http://www.ebi.ac.uk/ena/data/view/PRJEB6169). The metadata for these individuals is included in the BioSamples database under Project Identifier GSB-1451 (https://www.ebi.ac.uk/biosamples/groups/SAMEG317052). A small proportion of the tissues included in the Texel RNA-Seq dataset were not sampled in the BFxT gene expression atlas. Those unique to the Texel are largely drawn from the female reproductive, integument and nervous systems: cervix, corpus luteum, ovarian follicles, hypothalamus, brain stem, omentum and skin (side and back). Details of the Texel RNA-Seq libraries including tissue and cell type are included in S25 Table. The Texel samples were all prepared using the Illumina TruSeq stranded total RNA protocol with the Ribo-Zero Gold option for both cytoplasmic and mitochondrial rRNA removal, and sequenced using the Illumina HiSeq 2500 (151bp paired-end reads) [18]. As above, Kallisto was used to estimate expression level for all samples, using the revised reference transcriptome (from the ‘second pass’) as its index.

### Correcting for the Effect of Multiple Library Types

To correct for the confounding effect of multiple library types we applied a batch effect correction. We have previously validated this method using a subset of the sheep atlas expression atlas samples from BMDMs (+/- LPS) sequenced both as mRNA and total RNA libraries [31]. As described above, for the Kallisto second pass, we constrained the Kallisto index to contain only the transcripts of protein-coding genes, pseudogenes and processed pseudogenes, the majority of which are poly(A)+ and so are present in both mRNA-Seq and total RNA-Seq samples. We then calculated, per gene, the ratio of mean TPM across all mRNA-Seq libraries to mean TPM across all total RNA-Seq libraries. Given the scope of the tissues sampled for both library types (all major organ systems from both sexes and from different developmental states), neither mean is likely to be skewed by any tissue-specificity of expression. As such, any deviations of this ratio from 1 will reflect variance introduced by library type/depth. Thus, to correct each gene’s set of expression estimates for this effect of library type, we multiplied all total RNA-Seq TPMs by this ratio. To validate this approach we used principal component bi-plot analysis, described and shown in S1 Methods and S1 Fig.

### Gene Expression, Network Cluster Analysis and Annotation

Network cluster analysis of the sheep gene expression atlas was performed using Miru (Kajeka Ltd, Edinburgh, UK) [35-37]. In brief, similarities between individual gene expression profiles were determined by calculating a Pearson correlation matrix for both gene-to-gene and sample-to-sample comparisons, and filtering to remove relationships where *r* < 0.75. A network graph was constructed by connecting the remaining nodes (genes) with edges (where the correlation exceeded the threshold value). This graph was interpreted by applying the Markov Cluster algorithm (MCL) [38] at an inflation value (which determines cluster granularity) of 2.2. The local structure of the graph was then examined visually. Genes with robust co-expression patterns, implying related functions, clustered together, forming cliques of highly interconnected nodes. A principle of ‘guilt by association’ was then applied, i.e. the function of an unannotated gene could be inferred from the genes it clustered with [20, 136]. Expression profiles for each cluster were examined in detail to understand the significance of each cluster in the context of the biology of sheep tissues and cells. Clusters 1 to 50 (Table 2) were assigned a functional ‘class’ and ‘sub-class’ manually by first determining if multiple genes within a cluster shared a similar biological function based on both gene ontology [39], determined using the Bioconductor package ‘topGO’ [137] (GO term enrichment for clusters 1 to 50 is shown in S12 Table). We then compared the clusters with tissue- and cell-specific clusters in other large-scale network-based gene expression analyses including the pig gene expression atlas [6], the human protein atlas [69, 72, 138] and the mouse atlas [9, 139, 140]. More specific annotation of the GI tract clusters in sheep was based on network and pathway analysis from the sheep genome paper and a subsequent satellite publication [18, 21]. The gene component of all clusters can be found in S11 Table.

We assigned gene names to unnannotated genes in Oar v3.1 based on their co-expression pattern, tissue specificity, and reciprocal percent identity to a set of nine known ruminant proteomes (S7 Table). The annotation pipeline is described in detail in S1 Methods and included a set of quality categories summarised in S5 Table. We were able to assign gene names to >1000 previously unannotated genes in Oar v3.1. Candidate gene names are given as both a shortlist (S8 Table) and a longlist (S9 Table), the latter intended for manual review as informative annotations may still be made without every one of the above criteria being met.

## Abbreviations

AM, Alveolar Macrophage; BFxT, Scottish Blackface x Texel; BMDM, Bone Marrow Derived Macrophage; CNS, Central Nervous System; DE, Differential Expression; EBI, European Bioinformatics Institute; ENA, European Nucleotide Archive; FAANG, Functional Annotation of ANimal Genomes; GI, Gastrointestinal; HGNC, HUGO (Human Genome Organisation) Gene Nomenclature Committee; LPS, Lipopolysaccharide; MCL, Markov Cluster Algorithm; MCLi, Markov Cluster Algorithm Inflation; MDM, Monocyte Derived Macrophage; mRNA, messenger Ribonucleic Acid; NF-κB, Nuclear Factor Kappa-Light-Chain-Enhancer of Activated B-Cells; PBMC, Peripheral Blood Mononuclear Cell; RNA-Seq, RNA Sequencing; PPAR, Peroxisome Proliferator-Activated Receptor; rhCSF1, Recombinant Human Colony Stimulating Factor 1; SIGLEC, Sialic Acid-Binding Immunoglobulin-Like Lectin; SNP, Single Nucleotide Polymorphism

## Acknowledgments

We would like to thank the large number of people at the Roslin Institute who helped with the many aspects of the sheep atlas project. Douglas McGavin and David Chisholm coordinated the management of the sheep at Dryden Farm. Tim King, Peter Tennant, Adrian Ritchie, John Bracken and Pip Beard conducted post mortems on the animals. The lists of tissues for collection were developed by Erika Abbondati. Detailed dissection of the brain was performed by Fiona Houston. The tissue collection team included Heather Finlayson, Christine Burkhard, Alison Wilson, Ailsa Carlisle, Mark Barnett, Gemma Davis, Anna Raper, Rocio Rojo, Alex Brown, Chris Proudfoot, Laura Glendinning, Sara Clohisey, Yolanda Corripio-Miyar, Jack Ferguson, Karen Fernie and Jason Ioannidis. Blood Leukocytes were isolated by John Hopkins. RNA from the GI Tract was isolated by Gemma Davis and from embryonic fibroblasts by Charity Muriuki. Library preparation and sequencing was carried out by Edinburgh Genomics, The University of Edinburgh. We would also like to thank Norman Russell (The Roslin Institute) for generating the photographic images used and Andrew Su for allowing us to use BioGPS as the platform for visualistion of the expression data.

## Supplemental Material

**S1 Fig: Principal component analysis with all samples plotted in two dimensions using their projections onto the first two principal components.** For each sample, each gene’s expression level is taken as the mean TPM across all replicates, before (**A**) and after (**B**) any batch effect correction. Samples are coloured by organ system. Ellipses indicate confidence intervals of 95%. The shape of each point indicates each sample’s library type: mRNA-seq (circle) or total RNA-seq (triangle). Blastocyst samples are excluded for clarity as they are generated using a different experimental protocol. Before correction, points can be partitioned by shape (to the left and right of sub-Fig A), suggesting a batch effect – variation introduced by library type confounds variation by tissue type. After correction (sub-Fig B), there is no notable axis of variation that partitions points by shape – consequently, variation introduced by library type (a batch effect) does not confound variation by tissue type (which is biologically meaningful).

**S2 Fig: Sample-to-sample network graph analysis was used to validate the samples included in the sheep gene expression atlas dataset.** Each node represents a sample and each edge its connectivity to other samples in the dataset. A correlation of *r*=0.75 split the graph into 10 different clusters. The largest cluster (cluster 1) included the majority of samples (‘mixed tissues’), most of which were transcriptionally similar, while the remainder of the clusters comprised samples with distinctive transcriptional signatures such as macrophages (3) and abomasum. Spurious samples were easily identified if they were present in a cluster comprised of samples from a different tissue or cell type. Pearson Correlation *r*=0.75, MCLi = 2.2, nodes = 481 and edges = 23,903.

**S1 Methods:** Additional Methods

**S1 Dataset:** Gene Expression Level Atlas as transcripts per million (unaveraged)

**S2 Dataset:** Gene Expression Level Atlas as transcripts per million (averaged across biological replicates for each developmental stage)

**S3 Dataset:** Sex-biased Expression Atlas (based on expression estimates from 3 adult male and 3 adult female BFxT sheep)

**S1 Table:** Overview of animals used to generate the sheep atlas tissue subsets.

**S2 Table:** Details of library type and tissue/cell samples used in each subset of samples.

**S3 Table:** Number of reads, and number of aligned reads, per sample.

**S4 Table:** Genes undetected (TPM < 1) in every tissue/cell line of the expression atlas.

**S5 Table:** Quality categories for automated gene annotations.

**S6 Table:** Proportion of unannotated protein-coding genes assigned probable gene names.

**S7 Table:** Source of ruminant proteome data.

**S8 Table:** Candidate gene names for unannotated Oar v3.1 protein-coding genes: shortlist.

**S9 Table:** Candidate gene names for unannotated Oar v3.1 protein-coding genes: longlist.

**S10 Table:** Candidate gene descriptions for all unannotated Oar v3.1. protein-coding genes (potentially informative in the absence of a gene name).

**S11 Table:** Genes within each co-expression cluster.

**S12 Table:** GO term enrichment for co-expression clusters 1 to 50.

**S13 Table:** Manual annotation of cluster 15 (genes involved in oxidative phosphorylation).

**S14 Table:** Manual annotation of unannotated genes in cluster 12 (genes with a T-cell signature).

**S15 Table:** Manual annotation of unannotated genes in cluster 5 (genes with an alveolar macrophage signature).

**S16 Table:** Genes with five-fold sex-biased expression in at least one BFxT tissue.

**S17 Table:** GO term enrichment for the set of genes with five-fold sex-biased expression in at least one BFxT tissue.

**S18 Table:** Fold changes in expression level between BFxT and Texel sheep.

**S19 Table:** GO term enrichment for the set of genes differentially expressed between BFxT and Texel sheep.

**S20 Table:** Number of genes with detectable expression, per tissue.

**S21 Table:** Coding potential of putative novel CDS.

**S22 Table:** Number of genes with detectable expression, per gene type.

**S23 Table:** Proportion of Oar v3.1 gene, exon and transcript models in the StringTie assembly.

**S24 Table:** Novel transcript models in the StringTie assembly.

**S25 Table:** Details of the additional RNA-Seq libraries included from Texel sheep [18].

## References

1. Marino R, Atzori AS, D’Andrea M, Iovane G, Trabalza-Marinucci M, Rinaldi L. Climate change: Production performance, health issues, greenhouse gas emissions and mitigation strategies in sheep and goat farming. Small Ruminant Research. 2016;135:50-9. doi: doi.org/10.1016/j.smallrumres.2015.12.012.

2. Brito LF, Clarke SM, McEwan JC, Miller SP, Pickering NK, Bain WE, et al. Prediction of genomic breeding values for growth, carcass and meat quality traits in a multi-breed sheep population using a HD SNP chip. BMC Genetics. 2017;18(1):7. doi: 10.1186/s12863-017-0476-8.

3. Hayes BJ, Lewin HA, Goddard ME. The future of livestock breeding: genomic selection for efficiency, reduced emissions intensity, and adaptation. Trends in Genetics. 2013;29(4):206-14. doi: doi.org/10.1016/j.tig.2012.11.009.

4. Daetwyler HD, Hickey JM, Henshall JM, Dominik S, Gredler B, van der Werf JHJ, et al. Accuracy of estimated genomic breeding values for wool and meat traits in a multi-breed sheep population. Animal Production Science. 2010;50(12):1004-10. doi: doi.org/10.1071/AN10096.

5. Wickramasinghe S, Cánovas A, Rincón G, Medrano JF. RNA-Sequencing: A tool to explore new frontiers in animal genetics. Livestock Science. 2014;166:206-16. doi: doi.org/10.1016/j.livsci.2014.06.015.

6. Freeman TC, Ivens A, Baillie JK, Beraldi D, Barnett MW, Dorward D, et al. A gene expression atlas of the domestic pig. BMC Biology. 2012;10(1):90. doi: 10.1186/1741-7007-10-90.

7. Harhay GP, Smith TP, Alexander LJ, Haudenschild CD, Keele JW, Matukumalli LK, et al. An atlas of bovine gene expression reveals novel distinctive tissue characteristics and evidence for improving genome annotation. Genome Biology. 2010;11(10):R102. doi: 10.1186/gb-2010-11-10-r102.

8. Su AI, Cooke MP, Ching KA, Hakak Y, Walker JR, Wiltshire T, et al. Large-scale analysis of the human and mouse transcriptomes. Proc Natl Acad Sci USA. 2002;99. doi: 10.1073/pnas.012025199.

9. Su AI, Wiltshire T, Batalov S, Lapp H, Ching KA, Block D, et al. A gene atlas of the mouse and human protein-encoding transcriptomes. Proc Natl Acad Sci USA. 2004;101. doi: 1073/pnas.0400782101.

10. Mansour TA, Scott EY, Finno CJ, Bellone RR, Mienaltowski MJ, Penedo MC, et al. Tissue resolved, gene structure refined equine transcriptome. BMC Genomics. 2017;18(1):103. doi: 10.1186/s12864-016-3451-2.

11. Andersson R, Gebhard C, Miguel-Escalada I, Hoof I, Bornholdt J, Boyd M, et al. An atlas of active enhancers across human cell types and tissues. Nature. 2014;507(7493):455-61. doi: doi.org/10.1038/nature12787.

12. Lizio M, Harshbarger J, Shimoji H, Severin J, Kasukawa T, Sahin S, et al. Gateways to the FANTOM5 promoter level mammalian expression atlas. Genome Biology. 2015;16(1):22. doi: 10.1186/s13059-014-0560-6.

13. Forrest AR, Kawaji H, Rehli M, Baillie JK, De Hoon M, Haberle V, et al. A promoter-level mammalian expression atlas. Nature. 2014;507(7493):462-70. doi: doi.org/10.1038/nature13182.

14. Birney E, Stamatoyannopoulos JA, Dutta A, Guigo R, Gingeras TR, Margulies EH, et al. Identification and analysis of functional elements in 1% of the human genome by the ENCODE pilot project. Nature. 2007;447. doi: 10.1038/nature05874.

15. Melé M, Ferreira PG, Reverter F, DeLuca DS, Monlong J, Sammeth M, et al. The human transcriptome across tissues and individuals. Science. 2015;348(6235):660. doi: doi: 10.1126/science.aaa0355.

16. Bickhart DM, Rosen BD, Koren S, Sayre BL, Hastie AR, Chan S, et al. Single-molecule sequencing and chromatin conformation capture enable de novo reference assembly of the domestic goat genome. Nat Genet. 2017;49(4):643-50. doi: doi.org/10.1038/ng.3802.

17. Worley KC. A golden goat genome. Nat Genet. 2017;49(4):485-6. doi: 10.1038/ng.3824.

18. Jiang Y, Xie M, Chen W, Talbot R, Maddox JF, Faraut T, et al. The sheep genome illuminates biology of the rumen and lipid metabolism. Science. 2014;344(6188):1168. doi: doi: 10.1126/science.1252806.

19. Krupp M, Marquardt JU, Sahin U, Galle PR, Castle J, Teufel A. RNA-Seq Atlas - A reference database for gene expression profiling in normal tissue by next generation sequencing. Bioinformatics. 2012;28. doi: 10.1093/bioinformatics/bts084.

20. Oliver S. Proteomics: Guilt-by-association goes global. Nature. 2000;403(6770):601-3. doi: doi.org/10.1038/35001165.

21. Xiang R, Oddy VH, Archibald AL, Vercoe PE, Dalrymple BP. Epithelial, metabolic and innate immunity transcriptomic signatures differentiating the rumen from other sheep and mammalian gastrointestinal tract tissues. PeerJ. 2016;4:e1762. doi: 10.7717/peerj.1762.

22. Mabbott NA, Kenneth Baillie J, Hume DA, Freeman TC. Meta-analysis of lineage-specific gene expression signatures in mouse leukocyte populations. Immunobiology. 2010;215. doi: 10.1016/j.imbio.2010.05.012.

23. Mabbott NA, Kenneth Baillie J, Kobayashi A, Donaldson DS, Ohmori H, Yoon SO, et al. Expression of mesenchyme-specific gene signatures by follicular dendritic cells: insights from the meta-analysis of microarray data from multiple mouse cell populations. Immunology. 2011;133. doi: 10.1111/j.1365-2567.2011.03461.x.

24. Clop A, Marcq F, Takeda H, Pirottin D, Tordoir X, Bibe B, et al. A mutation creating a potential illegitimate microRNA target site in the myostatin gene affects muscularity in sheep. Nat Genet. 2006;38(7):813-8. doi: doi.org/10.1038/ng1810.

25. Blackface Sheep Breeders Association [1st March 2017]. Available from: http://www.scottish-blackface.co.uk/.

26. Andersson L, Archibald AL, Bottema CD, Brauning R, Burgess SC, Burt DW, et al. Coordinated international action to accelerate genome-to-phenome with FAANG, the Functional Annotation of Animal Genomes project. Genome Biology. 2015;16(1):57. doi: 10.1186/s13059-015-0622-4.

27. Tuggle CK, Giuffra E, White SN, Clarke L, Zhou H, Ross PJ, et al. GO-FAANG meeting: a Gathering On Functional Annotation of Animal Genomes. Animal Genetics. 2016;47(5):528-33. doi: 10.1111/age.12466.

28. Carninci P, Kasukawa T, Katayama S, Gough J, Frith MC, Maeda N, et al. The transcriptional landscape of the mammalian genome. Science. 2005;309. doi: 10.1126/science.1112014.

29. Bray NL, Pimentel H, Melsted P, Pachter L. Near-optimal probabilistic RNA-seq quantification. Nat Biotech. 2016;34(525–527). doi: doi.org/10.1038/nbt.3519.

30. Robert C, Watson M. Errors in RNA-Seq quantification affect genes of relevance to human disease. Genome Biology. 2015;16:177. doi: 10.1186/s13059-015-0734-x.

31. Bush SJ, McCulloch MEB, Summers KM, Hume DA, Clark EL. Integration of quantitated expression estimates from polyA-selected and rRNA-depleted RNA-seq libraries. BMC Bioinformatics. 2017;In Press.

32. Guo S, Lim D, Dong Z, Saunders TL, Ma PX, Marcelo CL, et al. Dentin sialophosphoprotein: a regulatory protein for dental pulp stem cell identity and fate. Stem Cells Dev. 2014;23(23):2883-94. doi: 10.1089/scd.2014.0066.

33. Wyatt K, Gao C, Tsai JY, Fariss RN, Ray S, Wistow G. A role for lengsin, a recruited enzyme, in terminal differentiation in the vertebrate lens. J Biol Chem. 2008;283(10):6607-15. doi: 10.1074/jbc.M709144200.

34. Pruitt KD, Tatusova T, Maglott DR. NCBI Reference Sequence (RefSeq): a curated non-redundant sequence database of genomes, transcripts and proteins. Nucleic acids research. 2005;33(Database Issue):D501-D4. doi: 10.1093/nar/gki025.

35. Freeman TC, Goldovsky L, Brosch M, van Dongen S, Maziere P, Grocock RJ, et al. Construction, visualisation, and clustering of transcription networks from microarray expression data. PLoS Computational Biology. 2007;3(10):2032-42. doi: 10.1371/journal.pcbi.0030206.

36. Theocharidis A, van Dongen S, Enright AJ, Freeman TC. Network visualization and analysis of gene expression data using BioLayout Express(3D). Nat Protoc. 2009;4(10):1535-50. Epub 2009/10/03. doi: 10.1038/nprot.2009.177.

37. Kajeka. Miru 2016 [16th March 2017]. Available from: https://kajeka.com/miru/miru-about/.

38. van Dongen S, Abreu-Goodger C. Using MCL to extract clusters from networks. Methods Mol Biol. 2012;804:281-95. doi: 10.1007/978-1-61779-361-5_15.

39. Ashburner M, Ball CA, Blake JA, Botstein D, Butler H, Cherry JM, et al. Gene ontology: tool for the unification of biology. The Gene Ontology Consortium. Nat Genet. 2000;25. doi: 10.1038/75556.

40. Hume DA, Summers KM, Raza S, Baillie JK, Freeman TC. Functional clustering and lineage markers: insights into cellular differentiation and gene function from large-scale microarray studies of purified primary cell populations. Genomics. 2010;95. doi: 10.1016/j.ygeno.2010.03.002.

41. Stumvoll M, Meyer C, Perriello G, Kreider M, Welle S, Gerich J. Human kidney and liver gluconeogenesis: evidence for organ substrate selectivity. American Journal of Physiology - Endocrinology And Metabolism. 1998;274(5):E817.

42. Ayyar VS, Almon RR, DuBois DC, Sukumaran S, Qu J, Jusko WJ. Functional proteomic analysis of corticosteroid pharmacodynamics in rat liver: Relationship to hepatic stress, signaling, energy regulation, and drug metabolism. Journal of Proteomics. doi: doi.org/10.1016/j.jprot.2017.03.007.

43. FANTOM Consortium. ZENBU: a collaborative, omics data integration and interactive visualization system 2017 [27th March 2017]. Available from: http://fantom.gsc.riken.jp/zenbu/.

44. Severin J, Lizio M, Harshbarger J, Kawaji H, Daub CO, Hayashizaki Y, et al. Interactive visualization and analysis of large-scale sequencing datasets using ZENBU. Nat Biotech. 2014;32(3):217-9. doi: doi.org/10.1038/nbt.2840.

45. Basten SG, Giles RH. Functional aspects of primary cilia in signaling, cell cycle and tumorigenesis. Cilia. 2013;2(1):6. doi: 10.1186/2046-2530-2-6.

46. Izawa I, Goto H, Kasahara K, Inagaki M. Current topics of functional links between primary cilia and cell cycle. Cilia. 2015;4(1):12. doi: 10.1186/s13630-015-0021-1.

47. Jackson PK. Do cilia put brakes on the cell cycle? Nat Cell Biol. 2011;13. doi: 10.1038/ncb0411-340.

48. Doig TN, Hume DA, Theocharidis T, Goodlad JR, Gregory CD, Freeman TC. Coexpression analysis of large cancer datasets provides insight into the cellular phenotypes of the tumour microenvironment. BMC Genomics. 2013;14(1):469. doi: 10.1186/1471-2164-14-469.

49. Calvo SE, Clauser KR, Mootha VK. MitoCarta2.0: an updated inventory of mammalian mitochondrial proteins. Nucleic Acids Research. 2016;44(Database issue):D1251-D7. doi: 10.1093/nar/gkv1003.

50. Sharma S, Sud N, Wiseman DA, Carter AL, Kumar S, Hou Y, et al. Altered carnitine homeostasis is associated with decreased mitochondrial function and altered nitric oxide signaling in lambs with pulmonary hypertension. American Journal of Physiology - Lung Cellular and Molecular Physiology. 2008;294(1):L46. doi: 10.1152/ajplung.00247.2007.

51. McCommis KS, Finck BN. Mitochondrial pyruvate transport: a historical perspective and future research directions. The Biochemical Journal. 2015;466(3):443-54. doi: 10.1042/BJ20141171.

52. Xiang R, McNally J, Rowe S, Jonker A, Pinares-Patino CS, Oddy VH, et al. Gene network analysis identifies rumen epithelial cell proliferation, differentiation and metabolic pathways perturbed by diet and correlated with methane production. Scientific Reports. 2016;6:39022. doi: doi.org/10.1038/srep39022.

53. Hofmann RR. Evolutionary steps of ecophysiological adaptation and diversification of ruminants: a comparative view of their digestive system. Oecologia. 1989;78(4):443-57. doi: 10.1007/BF00378733.

54. Rabbani I, Siegling-Vlitakis C, Noci B, Martens H. Evidence for NHE3-mediated Na transport in sheep and bovine forestomach. American Journal of Physiology - Regulatory, Integrative and Comparative Physiology. 2011;301(2):R313. doi: 10.1152/ajpregu.00580.2010.

55. Foster AM, Baliwag J, Chen CS, Guzman AM, Stoll SW, Gudjonsson JE, et al. IL-36 Promotes Myeloid Cell Infiltration, Activation, and Inflammatory Activity in Skin. The Journal of Immunology. 2014;192(12):6053. doi: 10.4049/jimmunol.1301481.

56. Perkins ND. Integrating cell-signalling pathways with NF-[kappa]B and IKK function. Nat Rev Mol Cell Biol. 2007;8(1):49-62. doi: 10.1038/nrm2083.

57. Palmer C, Diehn M, Alizadeh AA, Brown PO. Cell-type specific gene expression profiles of leukocytes in human peripheral blood. BMC Genomics. 2006;7(1):115. doi: 10.1186/1471-2164-7-115.

58. Margraf S, Garner LI, Wilson TJ, Brown MH. A polymorphism in a phosphotyrosine signalling motif of CD229 (Ly9, SLAMF3) alters SH2 domain binding and T-cell activation. Immunology. 2015;146(3):392-400. doi: 10.1111/imm.12513.

59. Akira S, Takeda K. Toll-like receptor signalling. Nat Rev Immunol. 2004;4(7):499-511. doi: 10.1038/nri1391.

60. Baillie JK, Arner E, Daub C, De Hoon M, Itoh M, Kawaji H, et al. Analysis of the human monocyte-derived macrophage transcriptome and response to lipopolysaccharide provides new insights into genetic aetiology of inflammatory bowel disease. PLoS Genetics. 2017;13(3):e1006641. doi: 10.1371/journal.pgen.1006641.

61. Pflanz S, Timans JC, Cheung J, Rosales R, Kanzler H, Gilbert J, et al. IL-27, a Heterodimeric Cytokine Composed of EBI3 and p28 Protein, Induces Proliferation of Naive CD4+ T Cells. Immunity. 2002;16(6):779-90. doi: doi.org/10.1016/S1074-7613(02)00324-2.

62. Hume DA, Ross IL, Himes SR, Sasmono RT, Wells CA, Ravasi T. The mononuclear phagocyte system revisited. Journal of Leukocyte Biology. 2002;72(4):621-7.

63. Moestrup SK, Moller HJ. CD163: a regulated hemoglobin scavenger receptor with a role in the anti-inflammatory response. Annals of Medicine. 2004;36(5):347-54.

64. Bain CC, Mowat AM. Intestinal macrophages – specialised adaptation to a unique environment. European Journal of Immunology. 2011;41(9):2494-8. doi: 10.1002/eji.201141714.

65. Vogt L, Schmitz N, Kurrer MO, Bauer M, Hinton HI, Behnke S, et al. VSIG4, a B7 family–related protein, is a negative regulator of T cell activation. Journal of Clinical Investigation. 2006;116(10):2817-26. doi: 10.1172/JCI25673.

66. Luo L, Bokil NJ, Wall AA, Kapetanovic R, Lansdaal NM, Marceline F, et al. SCIMP is a transmembrane non-TIR TLR adaptor that promotes proinflammatory cytokine production from macrophages. Nature Communications. 2017;8:14133. doi: 10.1038/ncomms14133.

67. Bonifer C, Hume DA. The transcriptional regulation of the Colony-Stimulating Factor 1 Receptor (csf1r) gene during hematopoiesis Frontiers in Bioscience. 2008;(13):549-60.

68. Hume DA, Yue X, Ross IL, Favot P, Lichanska A, Ostrowski MC. Regulation of CSF-1 receptor expression. Molecular Reproduction and Development. 1997;46(1):46-53. doi: 10.1002/(SICI)1098-2795(199701)46:1<46::AID-MRD8>3.0.CO;2-R.

69. Uhlén M, Fagerberg L, Hallström BM, Lindskog C, Oksvold P, Mardinoglu A, et al. Tissue-based map of the human proteome. Science. 2015;347(6220). doi: 10.1126/science.1260419.

70. Wu C, Jin X, Tsueng G, Afrasiabi C, Su AI. BioGPS: building your own mash-up of gene annotations and expression profiles. Nucleic Acids Research. 2016;44(D1):D313-D6. doi: 10.1093/nar/gkv1104.

71. Djureinovic D, Fagerberg L, Hallström B, Danielsson A, Lindskog C, Uhlén M, et al. The human testis-specific proteome defined by transcriptomics and antibody-based profiling. MHR: Basic science of reproductive medicine. 2014;20(6):476-88. doi: 10.1093/molehr/gau018.

72. Yu NY-L, Hallström BM, Fagerberg L, Ponten F, Kawaji H, Carninci P, et al. Complementing tissue characterization by integrating transcriptome profiling from the Human Protein Atlas and from the FANTOM5 consortium. Nucleic Acids Research. 2015;43(14):6787-98. doi: 10.1093/nar/gkv608.

73. Baillet A, Le Bouffant R, Volff JN, Luangpraseuth A, Poumerol E, Thépot D, et al. TOPAZ1, a Novel Germ Cell-Specific Expressed Gene Conserved during Evolution across Vertebrates. PLoS ONE. 2011;6(11):e26950. doi: 10.1371/journal.pone.0026950.

74. Olesen C, Larsen NJ, Byskov AG, Harboe TL, Tommerup N. Human FATE is a novel X-linked gene expressed in fetal and adult testis. Molecular and Cellular Endocrinology. 2001;184(1–2):25-32. doi: doi.org/10.1016/S0303-7207(01)00666-9.

75. Osaki E, Nishina Y, Inazawa J, Copeland NG, Gilbert DJ, Jenkins NA, et al. Identification of a novel Sry-related gene and its germ cell-specific expression. Nucleic Acids Research. 1999;27(12):2503-10.

76. Calloni R, Cordero EAA, Henriques JAP, Bonatto D. Reviewing and Updating the Major Molecular Markers for Stem Cells. Stem Cells and Development. 2013;22(9):1455-76. doi: 10.1089/scd.2012.0637.

77. Kehler J, Tolkunova E, Koschorz B, Pesce M, Gentile L, Boiani M, et al. Oct4 is required for primordial germ cell survival. EMBO reports. 2004;5(11):1078-83. doi: 10.1038/sj.embor.7400279.

78. Juengel JL, Bodensteiner KJ, Heath DA, Hudson NL, Moeller CL, Smith P, et al. Physiology of GDF9 and BMP15 signalling molecules. Animal Reproduction Science. 2004;82–83:447-60. doi: doi.org/10.1016/j.anireprosci.2004.04.021.

79. McNatty KP, Juengel JL, Wilson T, Galloway SM, Davis GH. Genetic mutations influencing ovulation rate in sheep. Reproduction, Fertility and Development. 2002;13(8):549-55.

80. Davis GH. Major genes affecting ovulation rate in sheep. Genetics Selection Evolution: GSE. 2005;37(Suppl 1):S11-S23. doi: 10.1186/1297-9686-37-S1-S11.

81. Palomera CL, Morales RAA. Genes with major effect on fertility in sheep. Revista Mexicana de Ciencias Pecuarias. 2014;5(1):107-30.

82. Davis GH. Fecundity genes in sheep. Animal Reproduction Science. 2004;82:247-53. doi: http://dx.doi.org/10.1016/j.anireprosci.2004.04.001.

83. Imahara SD, Jelacic S, Junker CE, O’Keefe GE. The influence of gender on human innate immunity. Surgery. 2005;138(2):275-82. doi: doi.org/10.1016/j.surg.2005.03.020.

84. Marriott I, Huet-Hudson YM. Sexual dimorphism in innate immune responses to infectious organisms. Immunologic Research. 2006;34(3):177-92. doi: 10.1385/IR:34:3:177.

85. Everhardt Queen A, Moerdyk-Schauwecker M, McKee LM, Leamy LJ, Huet YM. Differential Expression of Inflammatory Cytokines and Stress Genes in Male and Female Mice in Response to a Lipopolysaccharide Challenge. PLoS ONE. 2016;11(4):e0152289. doi: 10.1371/journal.pone.0152289.

86. Lamason R, Zhao P, Rawat R, Davis A, Hall JC, Chae JJ, et al. Sexual dimorphism in immune response genes as a function of puberty. BMC Immunology. 2006;7(1):2. doi: 10.1186/1471-2172-7-2.

87. Forde N, Maillo V, O’Gaora P, Simintiras CA, Sturmey RG, Ealy AD, et al. Sexually Dimorphic Gene Expression in Bovine Conceptuses at the Initiation of Implantation. Biology of Reproduction. 2016;95(4):92, 1-8-, 1-8. doi: 10.1095/biolreprod.116.139857.

88. Bermejo-Alvarez P, Rizos D, Rath D, Lonergan P, Gutierrez-Adan A. Sex determines the expression level of one third of the actively expressed genes in bovine blastocysts. Proc Natl Acad Sci USA. 2010;107. doi: 10.1073/pnas.0913843107.

89. Mentzel CMJ, Anthon C, Jacobsen MJ, Karlskov-Mortensen P, Bruun CS, Jørgensen CB, et al. Gender and Obesity Specific MicroRNA Expression in Adipose Tissue from Lean and Obese Pigs. PLoS ONE. 2015;10(7):e0131650. doi: 10.1371/journal.pone.0131650.

90. Zhang J, Zhou C, Ma J, Chen L, Jiang A, Zhu L, et al. Breed, sex and anatomical location-specific gene expression profiling of the porcine skeletal muscles. BMC Genetics. 2013;14(1):53. doi: 10.1186/1471-2156-14-53.

91. Wood Shona H, Christian Helen C, Miedzinska K, Saer Ben RC, Johnson M, Paton B, et al. Binary Switching of Calendar Cells in the Pituitary Defines the Phase of the Circannual Cycle in Mammals. Current Biology. 2015;25(20):2651-62. doi: doi.org/10.1016/j.cub.2015.09.014.

92. Bjelobaba I, Janjic MM, Kucka M, Stojilkovic SS. Cell Type-Specific Sexual Dimorphism in Rat Pituitary Gene Expression During Maturation. Biology of Reproduction. 2015;93(1):21. doi: 10.1095/biolreprod.115.129320.

93. Oidovsambuu O, Nyamsuren G, Liu S, Göring W, Engel W, Adham IM. Adhesion Protein VSIG1 Is Required for the Proper Differentiation of Glandular Gastric Epithelia. PLoS ONE. 2011;6(10):e25908. doi: 10.1371/journal.pone.0025908.

94. Ellegren H, Parsch J. The evolution of sex-biased genes and sex-biased gene expression. Nat Rev Genet. 2007;8(9):689-98. doi: 10.1038/nrg2167.

95. Gillespie JR, Flanders F. Modern Livestock & Poultry Production. 8th Edition ed: Delmar; 2009.

96. Zonabend König E, Ojango JMK, Audho J, Mirkena T, Strandberg E, Okeyo AM, et al. Live weight, conformation, carcass traits and economic values of ram lambs of Red Maasai and Dorper sheep and their crosses. Tropical Animal Health and Production. 2017;49(1):121-9. doi: 10.1007/s11250-016-1168-5.

97. Zonabend König E, Strandberg E, Ojango JMK, Mirkena T, Okeyo AM, Philipsson J. Purebreeding of Red Maasai and crossbreeding with Dorper sheep in different environments in Kenya. Journal of Animal Breeding and Genetics. 2017:n/a-n/a. doi: 10.1111/jbg.12260.

98. Rashid MM, Runci A, Polletta L, Carnevale I, Morgante E, Foglio E, et al. Muscle LIM protein/CSRP3: a mechanosensor with a role in autophagy. Cell Death Discovery. 2015;1:15014. doi: doi.org/10.1038/cddiscovery.2015.14.

99. Jan AT, Lee EJ, Choi I. Fibromodulin: A regulatory molecule maintaining cellular architecture for normal cellular function. Int J Biochem Cell Biol. 2016;80:66-70. Epub 2016/10/04. doi: 10.1016/j.biocel.2016.09.023.

100. Funderburgh JL, Mann MM, Funderburgh ML. Keratocyte Phenotype Mediates Proteoglycan Structure: A Role for Fibroblasts in Corneal Fibrosis. The Journal of Biological Chemistry. 2003;278(46):45629-37. doi: 10.1074/jbc.M303292200.

101. Hadler-Olsen E, Solli AI, Hafstad A, Winberg JO, Uhlin-Hansen L. Intracellular MMP-2 activity in skeletal muscle is associated with type II fibers. J Cell Physiol. 2015;230(1):160-9. Epub 2014/06/07. doi: 10.1002/jcp.24694.

102. Beard NA, Laver DR, Dulhunty AF. Calsequestrin and the calcium release channel of skeletal and cardiac muscle. Prog Biophys Mol Biol. 2004;85(1):33-69. Epub 2004/03/31. doi: 10.1016/j.pbiomolbio.2003.07.001.

103. Tellam RL, Cockett NE, Vuocolo T, Bidwell CA. Genes Contributing to Genetic Variation of Muscling in Sheep. Frontiers in Genetics. 2012;3:164. doi: 10.3389/fgene.2012.00164.

104. Miar Y, Salehi A, Kolbehdari D, Aleyasin SA. Application of myostatin in sheep breeding programs: A review. Molecular Biology Research Communications. 2014;3(1):33-43.

105. Cassar-Malek I, Passelaigue F, Bernard C, Léger J, Hocquette J-F. Target genes of myostatin loss-of-function in muscles of late bovine fetuses. BMC Genomics. 2007;8:63. doi: DOI: 10.1186/1471-2164-8-63.

106. Wang M, Yu H, Kim YS, Bidwell CA, Kuang S. Myostatin facilitates slow and inhibits fast myosin heavy chain expression during myogenic differentiation. Biochemical and Biophysical Research Communications. 2012;426(1):83-8. doi: doi.org/10.1016/j.bbrc.2012.08.040.

107. Allen DL, Loh AS. Posttranscriptional mechanisms involving microRNA-27a and b contribute to fast-specific and glucocorticoid-mediated myostatin expression in skeletal muscle. American Journal of Physiology - Cell Physiology. 2011;300(1):C124-C37. doi: 10.1152/ajpcell.00142.2010.

108. Mosca B, Eckhardt J, Bergamelli L, Treves S, Bongianino R, De Negri M, et al. Role of the JP45-Calsequestrin Complex on Calcium Entry in Slow Twitch Skeletal Muscles. Journal of Biological Chemistry. 2016;291(28):14555-65. doi: 10.1074/jbc.M115.709071.

109. Moran CM, Garriock RJ, Miller MK, Heimark RL, Mudry RE, Gregorio CC, et al. Expression of the fast twitch troponin complex, fTnT, fTnI and fTnC, in vascular smooth muscle. Cell motility and the cytoskeleton. 2008;65(8):652-61. doi: 10.1002/cm.20291.

110. Odermatt A, Becker S, Khanna VK, Kurzydlowski K, Leisner E, Pette D, et al. Sarcolipin Regulates the Activity of SERCA1, the Fast-twitch Skeletal Muscle Sarcoplasmic Reticulum Ca2+-ATPase. Journal of Biological Chemistry. 1998;273(20):12360-9. doi: 10.1074/jbc.273.20.12360.

111. Joo ST, Kim GD, Hwang YH, Ryu YC. Control of fresh meat quality through manipulation of muscle fiber characteristics. Meat Science. 2013;95(4):828-36. doi: doi.org/10.1016/j.meatsci.2013.04.044.

112. Goel M, Sienkiewicz AE, Picciani R, Wang J, Lee RK, Bhattacharya SK. Cochlin, Intraocular Pressure Regulation and Mechanosensing. PLoS ONE. 2012;7(4):e34309. doi: 10.1371/journal.pone.0034309.

113. Seidenbecher CI, Smalla KH, Fischer N, Gundelfinger ED, Kreutz MR. Brevican isoforms associate with neural membranes. J Neurochem. 2002;83(3):738-46.

114. Yamaguchi Y. Brevican: a major proteoglycan in adult brain. Perspect Dev Neurobiol. 1996;3(4):307-17.

115. Holz A, Schwab ME. Developmental expression of the myelin gene MOBP in the rat nervous system. J Neurocytol. 1997;26(7):467-77.

116. Yoshikawa F, Sato Y, Tohyama K, Akagi T, Hashikawa T, Nagakura-Takagi Y, et al. Opalin, a Transmembrane Sialylglycoprotein Located in the Central Nervous System Myelin Paranodal Loop Membrane. The Journal of Biological Chemistry. 2008;283(30):20830-40. doi: 10.1074/jbc.M801314200.

117. Boggs JM. Myelin basic protein: a multifunctional protein. Cellular and Molecular Life Sciences. 2006;63(17):1945-61. doi: 10.1007/s00018-006-6094-7.

118. Dwyer CM, Lawrence AB, Brown HE, Simm G. Effect of ewe and lamb genotype on gestation length, lambing ease and neonatal behaviour of lambs. Reprod Fertil Dev 1996;8(8):1123-9.

119. McCloskey E, McAdam JH. Grazing patterns and habitat selection of the Scottish Blackface compared with a crossbred, using GPS Satellite telemetry collars. Advances in Animal Biosciences. 2010;1(1):171. doi: 10.1017/S2040470010003146.

120. Wu C, MacLeod I, Su AI. BioGPS and MyGene.info: organizing online, gene-centric information. Nucleic Acids Research. 2013;41(D1):D561-D5. doi: 10.1093/nar/gks1114.

121. Wu C, Orozco C, Boyer J, Leglise M, Goodale J, Batalov S, et al. BioGPS: an extensible and customizable portal for querying and organizing gene annotation resources. Genome Biology. 2009;10(11):R130. doi: 10.1186/gb-2009-10-11-r130.

122. Kim D, Langmead B, Salzberg SL. HISAT: a fast spliced aligner with low memory requirements. Nat Meth. 2015;12(4):357-60. doi: doi.org/10.1038/nmeth.3317.

123. Pertea M, Pertea GM, Antonescu CM, Chang T-C, Mendell JT, Salzberg SL. StringTie enables improved reconstruction of a transcriptome from RNA-seq reads. Nat Biotech. 2015;33(3):290-5. doi: doi.org/10.1038/nbt.3122.

124. Miao X, Luo Q, Qin X. Genome-wide analysis reveals the differential regulations of mRNAs and miRNAs in Dorset and Small Tail Han sheep muscles. Gene. 2015;562(2):188-96. doi: doi.org/10.1016/j.gene.2015.02.070.

125. Suárez-Vega A, Gutiérrez-Gil B, Klopp C, Tosser-Klopp G, Arranz JJ. Variant discovery in the sheep milk transcriptome using RNA sequencing. BMC Genomics. 2017;18(1):170. doi: 10.1186/s12864-017-3581-1.

126. Sun L, Bai M, Xiang L, Zhang G, Ma W, Jiang H. Comparative transcriptome profiling of longissimus muscle tissues from Qianhua Mutton Merino and Small Tail Han sheep. Scientific Reports. 2016;6:33586. doi: doi.org/10.1038/srep33586.

127. Peñagaricano F, Valente BD, Steibel JP, Bates RO, Ernst CW, Khatib H, et al. Searching for causal networks involving latent variables in complex traits: Application to growth, carcass, and meat quality traits in pigs. Journal of Animal Science. 2015;93(10):4617-23. doi: 10.2527/jas.2015-9213.

128. Van Laere A-S, Nguyen M, Braunschweig M, Nezer C, Collette C, Moreau L, et al. A regulatory mutation in IGF2 causes a major QTL effect on muscle growth in the pig. Nature. 2003;425(6960):832-6. doi: doi.org/10.1038/nature02064.

129. Banos G, Bramis G, Bush SJ, Clark EL, McCulloch MEB, Smith J, et al. The genomic architecture of mastitis resistance in dairy sheep. 2017;Under Review.

130. Suárez-Vega A, Gutiérrez-Gil B, Klopp C, Tosser-Klopp G, Arranz J-J. Comprehensive RNA-Seq profiling to evaluate lactating sheep mammary gland transcriptome. Scientific Data. 2016;3:160051. doi: doi.org/10.1038/sdata.2016.51.

131. Kapetanovic R, Fairbairn L, Beraldi D, Sester DP, Archibald AL, Tuggle CK, et al. Pig bone marrow-derived macrophages resemble human macrophages in their response to bacterial lipopolysaccharide. J Immunol. 2012;188. doi: 10.4049/jimmunol.1102649.

132. Fairbairn L, Kapetanovic R, Beraldi D, Sester DP, Tuggle CK, Archibald AL, et al. Comparative Analysis of Monocyte Subsets in the Pig. The Journal of Immunology. 2013;190(12):6389-96. doi: 10.4049/jimmunol.1300365.

133. Montgomery GW, Sise JA. Extraction of DNA from sheep white blood cells. New Zeal and Journal of Agricultural Research. 1990;33(3):437-41. doi: 10.1080/00288233.1990.10428440.

134. Pervouchine DD, Djebali S, Breschi A, Davis CA, Barja PP, Dobin A, et al. Enhanced transcriptome maps from multiple mouse tissues reveal evolutionary constraint in gene expression. Nat Commun. 2015;6. doi: 10.1038/ncomms6903.

135. Chitwood J, Rincon G, Kaiser G, Medrano J, Ross P. RNA-seq analysis of single bovine blastocysts. BMC Genomics. 2013;14(1):350. doi: 10.1186/1471-2164-14-350.

136. Martherus RS, Sluiter W, Timmer ED, VanHerle SJ, Smeets HJ, Ayoubi TA. Functional annotation of heart enriched mitochondrial genes GBAS and CHCHD10 through guilt by association. Biochem Biophys Res Commun. 2010;402. doi: 10.1016/j.bbrc.2010.09.109.

137. Alexa A, Rahnenfuhrer J. topGO: Enrichment analysis for Gene Ontology 2010. Available from: http://www.bioconductor.org/packages/release/bioc/html/topGO.html.

138. Fagerberg L, Hallström BM, Oksvold P, Kampf C, Djureinovic D, Odeberg J, et al. Analysis of the Human Tissue-specific Expression by Genome-wide Integration of Transcriptomics and Antibody-based Proteomics. Molecular & Cellular Proteomics. 2014;13(2):397-406. doi: 10.1074/mcp.M113.035600.

139. Lein ES, Hawrylycz MJ, Ao N, Ayres M, Bensinger A, Bernard A, et al. Genome-wide atlas of gene expression in the adult mouse brain. Nature. 2007;445(7124):168-76. doi: doi.org/10.1038/nature05453.

140. Siddiqui AS, Khattra J, Delaney AD, Zhao Y, Astell C, Asano J, et al. A mouse atlas of gene expression: Large-scale digital gene-expression profiles from precisely defined developing C57BL/6J mouse tissues and cells. Proc Natl Acad Sci USA. 2005;102(51):18485-90. doi: 10.1073/pnas.0509455102.

141. Kumar S, Stecher G, Tamura K. MEGA7: Molecular Evolutionary Genetics Analysis Version 7.0 for Bigger Datasets. Molecular Biology and Evolution. 2016;33(7):1870-4. doi: 10.1093/molbev/msw054.

142. Rambaut A. FigTree v1.4.0 2016 [16th March 2017]. v1.4.3:[Available from: http://tree.bio.ed.ac.uk/software/figtree/.

